# Role of DEAD Box RNA helicases in low temperature adapted growth of Antarctic *Pseudomonas syringae* Lz4W

**DOI:** 10.1101/2022.10.22.513329

**Authors:** Ashaq Hussain, Malay Kumar Ray

## Abstract

Nucleic acid metabolism plays an important role in adaptation of organisms to environment. For adaptation to cold environment, organisms need to overcome the impediments in complex RNA secondary structures that are highly stabilized at low temperature affecting metabolism and growth of the organisms. Employing gene disruption through homologous recombination, we investigated the role of all major DEAD box RNA helicases (*rhlE, srmB, csdA, dbpA* and *rhlB*) in cold adapted growth of *Pseudomonas syringae*. Our findings concluded that *srmB* and *dbpA* genes are important for cold adapted growth, whereas *csdA* is indispensable for growth at low temperature. Helicase genes *rhlB* and *rhlE* have no appreciable role in cold adapted growth of *Pseudomonas syringae*. Functional complementation studies revealed that RNA helicases don’t have any redundant functions, as the roles performed by different helicases are individual and specific.

## Introduction

RNA helicases regulate cellular processes by modulating different aspects of RNA biology, from its synthesis to degradation, structural maturation (RNA processing) to the stability, and for interaction with metabolites to cellular proteins (RNA protein complexes) for sensing and responding to the cellular milieu[1, 2]. The DEAD-box RNA helicases constitute a specific group of RNA helicases, that derives its name from the highly conserved amino acid sequence in one of the motifs that reads as DEAD in single letter amino acid code[3]. The DEAD box (or slightly modified DExH/D box) proteins belong to the SF2 superfamily of RNA helicases which bind to ATP and RNA by two RecA like domains located on 350 to 400 residues long ‘helicase core” and perform the ATP (ATPase) dependent functions [4, 5]. These functions include canonical RNA duplex unwinding, RNA strand annealing, RNA folding (RNA chaperone), strand exchange, and protein displacement activity [1, 6, 7]. The conserved ‘helicase core’ of the RNA helicases is generally flanked by variable regions which are important for determining specificity of the enzyme [2]. The cellular ability to perform these functions at low temperature is critical for the cold adapted organisms, as RNA secondary structures are stabilized at lower temperatures [8].

In bacteria only a small number of genes (four to six) code for DEAD box proteins [9]. Gram-negative bacterium, such as *E. coli* contains five genes for different DEAD box RNA helicases while the Gram-positive bacterium such as *B. subtilis* contains four DEAD-box RNA helicase genes in the genome [7, 10, 11]. These helicases are involved in ribosome biogenesis, mRNA processing and translation. Many of these proteins are non-essential in the organisms at optimum temperature of growth, but variably important for growth at low temperature. In *E. coli*, none of the five DEAD box proteins (RhlE, SrmB, CsdA, DbpA and RhlB) are essential for growth at 37°C [12]. However, SrmB and CsdA are critical at low temperature (15-20°C) in contrast, RhlE, RhlB and DbpA are dispensable for growth at low-temperature [7, 8, 12, 13].

RhlE has been found associated with 70S ribosomes under normal conditions of growth. Its association to the ribosomal subunits (50S and 30S), however, has been proposed based on the analysis of the defective 40S particle in Δ*csdA*, Δ*srmB* and Δ*csdA*Δ*srmB* double mutant suggesting that the helicase may have a direct role in the ribosome biogenesis [7, 12, 13]. RhlE has been implicated in biogenesis and assembly of the 50S ribosomal subunit where it may be involved in interconversion of two different conformational structures of 23S rRNA specific for SrmB and DbpA [13]. It has been proposed that under the overexpressing conditions RhlE favors the SrmB specific structural conformation of 23S rRNA while it favors the CsdA specific conformation of 23S rRNA in the absence of RhlE. The possibility of the existence of two different conformations of 23S rRNA and their conversion explains the opposing effects of rhlE disruption in Δ*srmB* and Δ*csdA* genetic backgrounds [12, 13].

SrmB has been classified as a ribosome assembly factor [8, 14–16]. Disruption of *srmB* in *E. coli* has no associated phenotype at the optimum temperature (37°C) of growth, however, at 25°C the Δ*srmB* mutant displays a cold-sensitive phenotype [12, 15]. Molecular analysis indicated that the mutant cells exhibit higher amount of precursor 23S rRNAs that contain unprocessed nucleotides (7-9 bases) at 3’ ends and unprocessed 5’ ends (3-7 bases). Ribosome profiling revealed that the mutant cells contain reduced amount of 50S subunits and an accumulation of defective 40S particles [12, 15].

DEAD box helicase CsdA (also known as *DeaD* in *E. coli*) was found to be crucial for the growth of *E. coli* at low temperature[12]. Deletion of *csdA* gene (Δ*csdA* or Δ*deaD*) led not only to severe defects in growth and viability at the low temperature (20-25°C) but also to a slow growing phenotype at the optimum temperature (37°C). Δ*csdA* exhibited similar type of rRNA processing and ribosome assembly defects as shown by Δ*srmB* but to a higher degree, both at the low (25°C) and optimum (37°C) temperatures of growth [12, 17, 18]. The accumulation of defective ribosomal RNAs was 4 folds higher in Δ*csdA* than in wild-type [12]. This conclusion is consistent with the growth measurements that revealed at 37°C; each of the multiple helicase deletion strains (e.g., quadruple Δ*dbpA*Δ*rhlB*Δ*rhlE*Δ*srmB* mutants and quintuple Δ*csdA*Δ*dbpA*Δ*rhlB*Δ*rhlE*Δ*srmB* (Δ5) mutant grew more slowly than a wild-type strain but none of them grew significantly slower than the singly mutated Δ*csdA* strain [12].

Deletion of *rhlB* and *dbpA* helicases do not produce any temperature-sensitive phenotypes in *E. coli* However, a highly specific interaction of DbpA with 23S rRNA via its C-terminal domain points towards a role of the protein in ribosome biogenesis where the helicase acts possibly by structural rearrangement of the 23S rRNA [18–20]. Evidence for the role of DbpA in late stages of 50S subunit assembly was recently confirmed when *E. coli* strain over-expressing an inactive DbpA helicase displayed the cold-sensitive phenotype [20, 21]. Some of the late ribosome binding proteins that bind close to the DbpA biding site on 23S rRNA were absent in the ribosomes [22]

In *E. coli*, the mutant strains deleted for all five DEAD-box RNA helicases in different combinations showed that the strains with multiple knockouts grow slower than the wild-type at 37°C, but not significantly less than the single Δ*csdA* mutant indicating the important role of *csdA* in growth at all temperatures[12]. However, at low temperature (25°C) the growth analysis revealed that there is an increase in generation time with each successive gene deletion as a result of which the strain with all five gene deletions, strain (Δ5), displayed the longest generation time (181 hrs) as compared to the wild-type (72 hrs) and even Δ*csdA* (140 hrs) [12]. Analysis of the rRNAs was also in conformity with five-fold more accumulation of defective rRNAs in the Δ5 strain compared to wild-type. Ribosome profiling revealed a decrease in the 70S fraction and increase in the defective 40S fraction for all cold-sensitive mutant strains (Δ5, Δ*srmB*Δ*csdA*, and Δ*csdA*). The defect was severe in Δ5 strain, in which ribosome profile was more aberrant than in Δ*srmB*, Δ*csdA*, and Δ*dbpA* indicating a role of RhlE and RhlB helicases too in the ribosome biogenesis[12]. The growth analysis of the helicase mutants in minimal media showed some interesting differences [12]. At optimum temperature (37°C) there was no difference in the generation time between WT and any of the helicase mutant/s in the minimal growth medium; however, at 25°C, generation time of the Δ5 strain, although longer, was 5 times lesser, compared to the wild-type in rich medium [8, 12]

To understand the molecular mechanism of cold-adaptation our laboratory has been using the Antarctic bacterium *Pseudomonas syringae* Lz4W as model organism [23, 24]. The major findings from our laboratory have established that the stability of DNA and RNA secondary structures have played a critical role in the adaptation and evolution of DNA and RNA metabolic enzymes allowing the bacterial growth at low temperature. Any defect in these metabolic enzymes leads to cold sensitivity of the bacterium. For example, the inactivation of the exoribonuclease enzyme RNase R, a component of RNA degrading machinery in *P. syringae*, leads to lethality of the bacterium at cold, and the bacterium fails to grow at 4°C [25]. Similarly, inactivation of the DNA repairing RecBCD machinery leads to loss of chromosomal integrity, cell lethality, and hence the growth defect at low temperature [26, 27]. As a follow up of these basic findings we have addressed the issues further by investigating the importance of the DEAD box RNA helicases in cold-adaptation, as this group of enzymes play a major role in modulating the RNA secondary structures and RNA metabolism at low temperature.

Since DEAD box RNA helicases act by modulating the RNA secondary structures known to be stabilized at low temperatures to regulate different aspects of RNA biology and cell physiology of cold-adapted organisms, we took a genomic approach to identify and assess the different RNA helicases that are present in the genome of the Antarctic *P. syringae* Lz4W and needed for its psychrophilic adaptation. In *P. syringae* we identified genes for five major DEAD box RNA helicases (*rhlE, srmB, csdA, dbpA* and *rhlB*) encoding RhlE, SrmB, CsdA, DbpA and RhlB, respectively, that are present in most Gram-negative bacteria. All of these RNA helicases contain a set of highly conserved (Twelve) motifs including the ‘DEAD’ box (motif II) of the SF2 helicase superfamily [7, 28, 29]. Since most of the studies on DEAD-box RNA helicases so far have been carried out with the above five major DEAD-box RNA helicases of mesophilic *E. coli* [12], and *Bacillus Subtilis* [11], we also focused only on these five RNA helicases of the psychrophilic *P. syringae* for their role in cold-adaptation. Interestingly, we found that *Pseudomonas* and related bacteria contain a second gene homologue (*rhlE*-S) for the RhlE helicase which is shorter in length (445 residues, and named RhlE-S), which is different from the degradosome associated RhlE (618 residues).

Although the individual role of the DEAD-box RNA helicases can be inferred from the analysis of single deletion strains, physiologically the helicase genes work together, either co-operatively in combinations or individually by division of labor, within the cells. Sometimes, the effects of single gene inactivation might not show any phenotypic effect due to functional redundancy. Therefore, the question arises whether the cellular defects will be more pronounced if more than one helicase genes are successively deleted in different combination, leading to either additive effects or synergistic effects depending on their cellular interactions. To address this issue, we constructed several double-deletion strains and one triple-deletion mutant for the helicase genes in this study and assessed their growth characteristics at low (4°C) and optimum (22°C) temperatures.

Main objectives of the current study are (a) Identification and organization of DEAD box RNA helicase genes in *P. Syringae* genome (c) Sequence analysis of RNA helicase genes including the 5’ and 3’ UTR regions (d) To study the role of DEAD box RNA helicases in cold adapted growth of *P. syringae* and (e) Identification of any redundant/overlapping functions among helicases by performing functional complementation assay.

## Material and methods

### Bacterial cultures and growth media

The psychrophilic *P. syringae* Lz4W was routinely grown at 22°C or 4°C (for optimum and low temperatures respectively) in Antarctic bacterial medium (ABM) composed of 5 g/l peptone and 2.0 g/l yeast extract, as described earlier [26, 30]. *E. coli* strains were cultured at 37°C in Luria-Bertani (LB) medium, which contained 10g/l tryptone, 5 g/l yeast extract and 10 g/l NaCl [31]. For solid media, 15 g/l bacto-agar (Hi Media) was added to ABM or LB. Both ABM and LB media were supplemented with ampicillin (100μg/ml), kanamycin (50μg/ml), spectinomycin (100 μg/ml), gentamicin (15 μg/ml) and tetracycline (10 μg/ml) as per requirement.

Generation times were calculated from the growth curves of the different recombinant strains. Fresh ABM broth was inoculated with 1% of primary culture and incubated at 22C or 4C with constant shaking. Optical density of bacterial culture was measured after different time intervals at 600 nm [OD_600_] and plotted against time.

### Molecular biology methods

General molecular biology techniques including isolation of genomic DNA, polymerase chain reactions (PCR), restriction enzyme digestion and ligation, transformation etc. were performed as described [33]. All restriction enzymes, T4 DNA ligase and other enzymes used in this study were from New England Biolabs (NEB, USA). Polymerase chain reactions for gene amplification and site directed mutagenesis were carried out using high fidelity proof reading pfx DNA polymerase from Invitrogen (USA). Preparation of plasmid and purification of PCR products was done by designated Qiagen kits (Qiagen, Germany). Oligonucleotides were purchased from a commercial source (Bioserve Biotechnology, India). The conjugal transfer of recombinant plasmid into *P. syringae* was carried out by a biparental mating method using the donor *E. coli* strain S17-1, as described in [32].

### Live Dead staining and cell viability assay

Live/dead staining of *P. Syringae* cultures was performed by using LIVE/DEAD Bacterial BacLight Viability Kit [Invitrogen]. All the steps of sample preparation and staining were performed as directed by manufacturer[33].

### Construction of recombinant plasmids

All gene cloning experiments were performed in DH5α cells. The detailed methodology has been described previously [31, 34, 35]. Plasmids used for expression of RNA helicase proteins, genetic complementation of helicase mutants and disruption of helicase genes are listed in Table S1 to S3.

### Site directed mutagenesis

All site-specific mutations were introduced by using QuikChange Site-Directed Mutagenesis Kit [Agilent technologies, USA] according to the manufacturer’s instructions. Oligos used for insertion of mutations in a desired gene are listed in table S4.

### Generation of DEAD box RNA helicase mutant strains of *P. syringae* Lz4W

Disruption of target gene was achieved by gene replacement method using homologous recombination between plasmid born antibiotic cassette disrupted gene and chromosomal gene as reported earlier [25, 27]. A common method was followed for gene disruption, in which a selective marker (antibiotic resistance cassette) was inserted into the middle portion of target gene cloned in suicidal plasmid pJQ200SK [36] [26, 27, 37, 38]. For double crossover recombination to occur between the antibiotic cassette disrupted gene in suicidal plasmid and *P. syringae* chromosomal gene, approximately 400 to 500 base pairs of homologous DNA sequence were provided on either side of the antibiotic resistance cassette. Schematic representation for generation of suicidal plasmid constructs used for disruption of RNA helicase genes is shown in Fig. 5. Numbers above the gene(s) in schematic refer to nucleotide number of gene. Restriction sites used for cloning/insertion of antibiotic cassette are marked. The suicidal plasmid constructs generated in this study are pJQ*rhlE*-tet, pJQ*srmB*-kan, PJQ*csdA*-spec, pJQ*dbpA*-spec and pJQ*rhlB*-spec (Table. S3). oligonucleotides used for validation of gene disruptions in *P. syringae* are listed in table S5. Different RNA helicase mutant strains generated in this study are listed in table 1.

**Table 1.** Bacterial strains used for general cloning, protein expression and study of *P. Syringae* strains disrupted for different DEAD box RNA helicases.

| Bacterial strains | Genotype | Reference or source |
| --- | --- | --- |
| DH5α | F- ϕ80*lacZ*ΔM15 $\Delta$(*lacZYA-argF*) U169 *recA1 endA1 hsdR17* ($r_k^-$, $m_k^+$) *phoA supE44* λ- *thi-1 gyrA96 relA1* Used for all gene cloning purpose. | [35] |
| S17-1 | *F _ pro recA1 (r_ m_) RP4-2 integrated (Tc::Mu) (Km::Tn7) [Smr Tpr]* ; used as a donor strain in conjugation | [32] |
| BL21 (DE3) | *F^-ompT gal dcm lon hsdS<sub>B</sub>(r<sub>B</sub>^- m<sub>B</sub>^-) $\lambda$(DE3 [*lacI lacUV5-T7 gene 1 ind1 sam7 nin5*])*, Used for over expression of proteins under IPTG induction. | [39] |
| *P.syringae* (Lz4W) | Lz4W Amp<sup>r</sup>, Wild-type/Natural isolate | [23] |
| $\Delta rhIE$ | *rhIE:: ter<sup>r</sup>*<br>*P. Syringae* strain with disrupted *rhIE* gene. | This study |
| $\Delta srmB$ | *srmB:: kan<sup>r</sup>*<br>*P. Syringae* strain with disrupted *srmB* gene. | This study |
| $\Delta srmB\Delta rhIE$ | *srmB:: kan<sup>r</sup>, rhIE:: ter<sup>r</sup>*<br>*P. Syringae* strain with disrupted *srmB* and *rhIE* genes. | This study |
| $\Delta csdA$ | *csdA:: spec<sup>r</sup>*<br>*P. Syringae* strain with disrupted *csdA* gene. | This study |
| $\Delta csdA\Delta rhIE$ | *csdA:: spec<sup>r</sup>, rhIE:: ter<sup>r</sup>*<br>*P. Syringae* strain with disrupted *csdA* and *rhIE* genes. | This study |
| $\Delta dbpA$ | *dbpA:: spec<sup>r</sup>*<br>*P. Syringae* strain with disrupted *dbpA* and *rhIE* genes. | This study |
| $\Delta dbpA\Delta rhIE$ | *dbpA:: spec<sup>r</sup>, rhIE:: ter<sup>r</sup>*<br>*P. Syringae* strain with disrupted *dbpA* and *rhIE* genes. | This study |
| $\Delta dbpA\Delta srmB$ | *dbpA:: spec<sup>r</sup>, srmB:: kan<sup>r</sup>*<br>*P. Syringae* strain with disrupted *dbpA* and *srmB* genes. | This study |
| $\Delta rhIB$ | *rhIB:: spec<sup>r</sup>*<br>*P. Syringae* strain with disrupted *rhIB* gene. | This study |
| $\Delta rhIB\Delta rhIE$ | *rhIB:: spec<sup>r</sup>, rhIE:: ter<sup>r</sup>*<br>*P. Syringae* strain with disrupted *rhIB* and *rhIE* genes. | This study |
| $\Delta rhIB\Delta srmB$ | *rhIB:: spec<sup>r</sup> srmB:: kan<sup>r</sup>*<br>*P. Syringae* strain with disrupted *rhIB* and *srmB* genes. | This study |
| $\Delta rhIB\Delta rhIE\Delta srmB$ | *rhIB:: spec<sup>r</sup>, rhIE:: ter<sup>r</sup> srmB:: kan<sup>r</sup>*<br>*P. Syringae* strain with disrupted *rhIB*, *rhIE* and *srmB* genes. | This study |

### Construction of plasmids for expression and complementation Studies

The helicase genes cloned in pET28-*rhlE*, pET28-*srmB*, pET28-*csdA*, pET28-*dbpA* and pET28-*rhlB* [Table S1] were released along with the ribosome binding site (RBS) of pET28a with XbaI and SacI restriction enzymes, and cloned into the *XbaI/SacI* sites of broad host range plasmid pGL10 [39, 40]. The resultant plasmids *pGrhlE, pGsrmB, pGcsdA, pGdbpA* and *pGrhlB* (Table S5) harbouring the respective genes for RhlE, SrmB, CsdA, DbpA and RhlB helicases were propagated in *E. coli* DH5α and later transformed into *E. coli* S17-1 for their mobilization into *P. syringae* strains. The plasmids pGL10, pG*rhlE*, pG*srmB*, pG*csdA*, pG*dbpA* and pG*rhlB* were mobilized separately into each individual RNA helicase mutant of *P. syringae* by conjugation with *E. coli* S17-1[41]. All the strains used in genetic complementation study are listed in Table 2.

**Table 2:** *P. Syringae* strains used in functional complementation study of RNA helicase mutants.

| Plasmid | Description | Reference/<br>Source |
| --- | --- | --- |
| Lz4W (pGL10) | Wild type strain complemented by empty pGL10 plasmid. | This study |
| Lz4W (pGL10*rhlE*) | Wild type strain complemented by *rhlE*. | This study |
| Lz4W(pGL10*srmB*) | Wild type strain complemented by *srmB*. | This study |
| Lz4W (pGL10*csdA*) | Wild type strain complemented by *csdA*. | This study |
| Lz4W(pGL10*dbpA*) | Wild type strain complemented by *dbpA*. | This study |
| Lz4W (pGL10*rhlB*) | Wild type strain complemented by *rhlB*. | This study |
| Δ*rhlE*(pGL10) | Δ*rhlE* strain complemented by empty pGL10 plasmid. | This study |
| Δ*rhlE*(pGL10*rhlE*) | Δ*rhlE* strain complemented by *rhlE*. | This study |
| Δ*rhlE*(pGL10*srmB*) | Δ*rhlE* strain complemented by *srmB*. | This study |
| Δ*rhlE*(pGL10*csdA*) | Δ*rhlE* strain complemented by *csdA*. | This study |
| Δ*rhlE*(pGL10*dbpA*) | Δ*rhlE* strain complemented by *dbpA*. | This study |
| Δ*rhlE*(pGL10*rhlB*) | Δ*rhlE* strain complemented by *rhlB*. | This study |
| Δ*srmB*(pGL10) | Δ*srmB* strain complemented by empty pGL10 plasmid. | This study |
| Δ*srmB*(pGL10*rhlE*) | Δ*srmB* strain complemented by *rhlE*. | This study |
| Δ*srmB*(pGL10*srmB*) | Δ*srmB* strain complemented by *srmB*. | This study |
| Δ*srmB*(pGL10*csdA*) | Δ*srmB* strain complemented by *csdA*. | This study |
| Δ*srmB*(pGL10*dbpA*) | Δ*srmB* strain complemented by *dbpA*. | This study |
| Δ*srmB*(pGL10*rhlB*) | Δ*srmB* strain complemented by *rhlB*. | This study |
| Δ*csdA*(pGL10) | Δ*csdA* strain complemented by empty pGL10 plasmid. | This study |
| Δ*csdA*(pGL10*rhlE*) | Δ*csdA* strain complemented by *rhlE*. | This study |
| Δ*csdA*(pGL10*srmB*) | Δ*csdA* strain complemented by *srmB*. | This study |
| Δ*csdA*(pGL10*csdA*) | Δ*csdA* strain complemented by *csdA*. | This study |
| Δ*csdA*(pGL10*dbpA*) | Δ*csdA* strain complemented by *dbpA*. | This study |
| Δ*csdA*(pGL10*rhlB*) | Δ*csdA* strain complemented by *rhlB*. | This study |
| Δ*dbpA*(pGL10) | Δ*dbpA* strain complemented by empty pGL10 plasmid. | This study |
| Δ*dbpA*(pGL10*rhlE*) | Δ*dbpA* strain complemented by *rhlE*. | This study |
| Δ*dbpA*(pGL10*srmB*) | Δ*dbpA* strain complemented by *srmB*. | This study |
| Δ*dbpA*(pGL10*csdA*) | Δ*dbpA* strain complemented by *csdA*. | This study |
| Δ*dbpA*(pGL10*dbpA*) | Δ*dbpA* strain complemented by *dbpA*. | This study |
| Δ*dbpA*(pGL10*rhlB*) | Δ*dbpA* strain complemented by *rhlB*. | This study |
| Δ*rhlB*(pGL10) | Δ*rhlB* strain complemented by empty pGL10 plasmid. | This study |
| Δ*rhlB*(pGL10*rhlE*) | Δ*rhlB* strain complemented by *rhlE*. | This study |
| Δ*rhlB*(pGL10*rhlE*) | Δ*rhlB* strain complemented by *rhlE*. | This study |
| Δ*rhlB*(pGL10*srmB*) | Δ*rhlB* strain complemented by *srmB*. | This study |
| Δ*rhlB*(pGL10*csdA*) | Δ*rhlB* strain complemented by *csdA*. | This study |
| Δ*rhlB*(pGL10*dbpA*) | Δ*rhlB* strain complemented by *dbpA*. | This study |
| Δ*rhlB*(pGL10*rhlB*) | Δ*rhlB* strain complemented by *rhlB*. | This study |

## Results

### Genome organization of the DEAD box RNA helicase genes in *P. syringae*

Analysis of the *P. syringae* Lz4W genome sequence helped us to identify six DEAD-box RNA helicase genes in the chromosome of the bacterium. We identified the genes (*rhlE*, *srmB*, *csdA*, *dbpA* and *rhlB*) for the five major DEAD-box RNA helicases of Gram-negative bacteria which encoded RhlE, SrmB, CsdA, DbpA and RhlB, respectively. An additional helicase gene which is a shorter homologue of *rhlE*, named *rhlE-S* encoding the 445 residues RhlE-S as compared to the degradosome associated larger RhlE (618 residues) was also found in the genome. The organization of DEAD-box RNA helicase genes namely *rhlE, rhlE-S, srmB, csdA, dbpA* and *rhlB* encoded by genetic loci, B195_4530, B195_12841, B195_08192, B195_19396, B195_14360 and B195_05446 respectively, on the *P. syringae* genome are shown in Fig. 1. From the analysis of directions of transcriptions and the distance between the genes it was inferred that these RNA helicase genes were monocistronic and dispersed over different parts of the *P. syringae* genome.

**Fig. 1.**
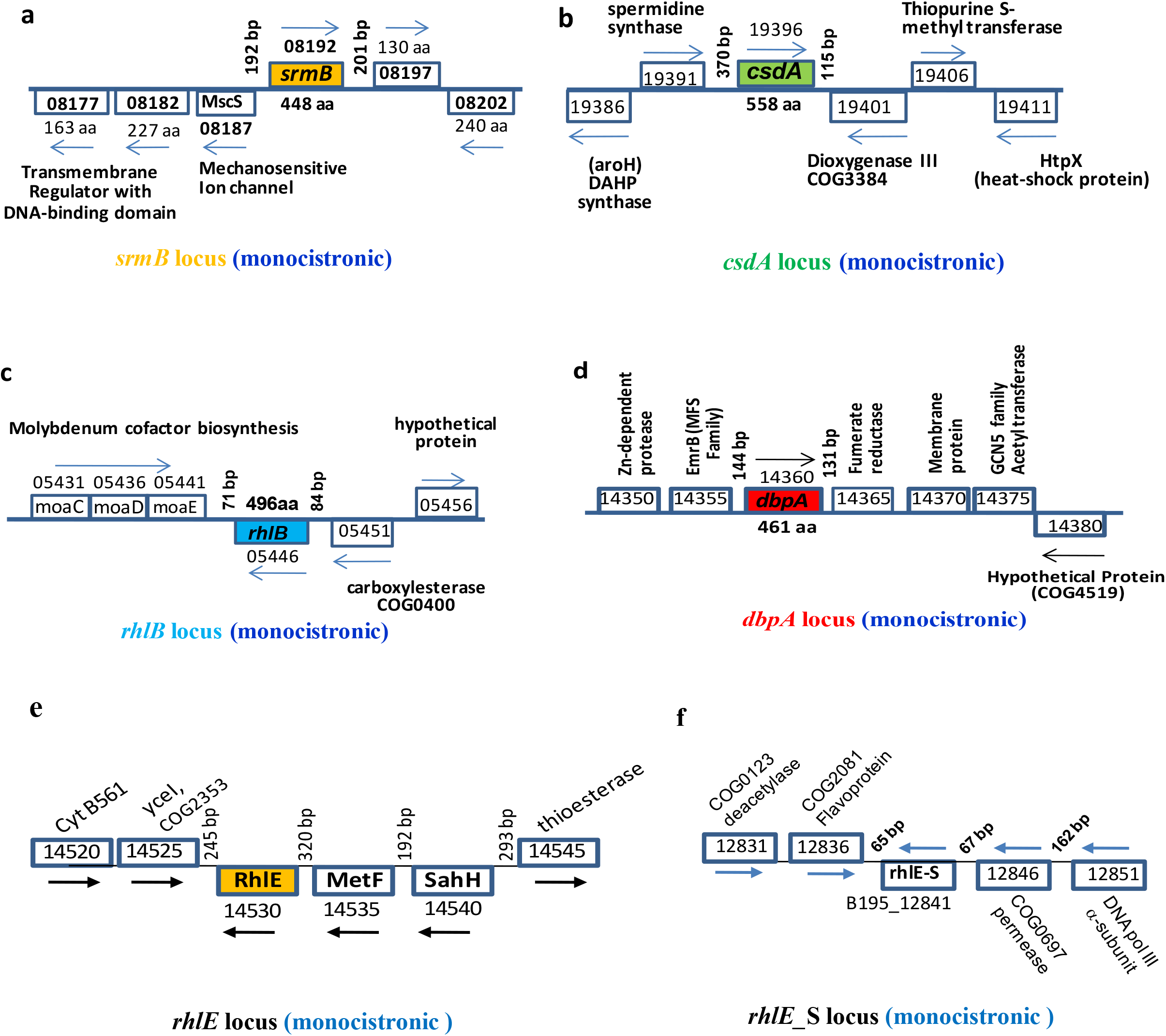
Genomic organization of the five DEAD-box RNA helicase genes of *P. syringae* Lz4W. The helicase genes and their surrounding upstream and downstream genes have been indicated by their respective locus tag numbers in the *P. syringae* genome. The locus tags in the annotated genome (acc no. AOGS01000001 to 42) at the NCBI site starts with B195_xxxxx numbers. For clarity on the Figure, the common prefix ‘B195_‘ has been omitted in the diagram. The direction of transcription has been shown by arrows on above or below the gene-ORF boxes. The encoded gene product lengths has been shown by amino acid numbers. (**a**) *srmB* gene locus, (**b**) *csdA* gene locus, (**c**) *rhlB* locus, (**d**) *dbpA* gene locus, (**e**) *rhlE* gene locus **(f).**rhlE_S gene locus.

The upstream and downstream intergenic spacers contained the putative promoter and regulatory sequences of the helicase genes which were of variable lengths (Fig. 2). The *rhlB* upstream and downstream intergenic spacers had the shortest lengths corresponding to 84 bp and 71 bp sequences, respectively. The *csdA* had the longest upstream intergenic spacer spanning 370 bp sequences. The *srmB* and *dbpA* genes had modest upstream intergenic spacers of 192 bp and 144 bp, respectively. The downstream intergenic distances were also variable in length which contained the putative transcription terminators for the genes (Fig. 2a).

**Fig. 2.**
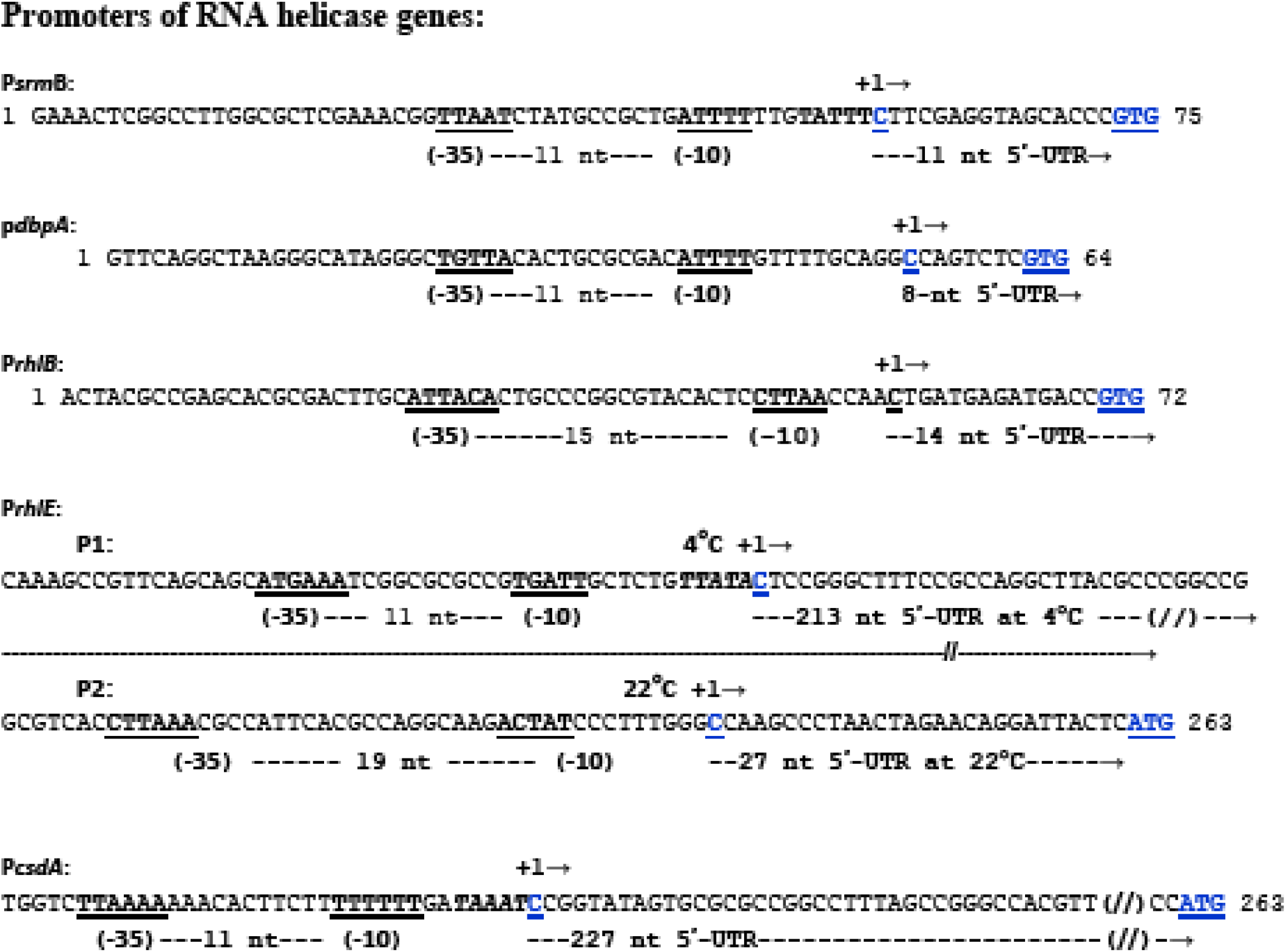
Putative regulatory regions of *P. syringae* helicase genes showing promoter characteristics and 5’-UTR lengths of the five major DEAD-box RNA helicase genes of *P. syringae* Lz4W. The ‘-10’and ‘35’ sequences of the promoters have been underlined at the upstream of (+1) transcription-start sites that are all located ‘ATG’ or GTG’ translation initiation codons of the respective genes. The lengths of the 5’-UTR have been indicated for each of the transcripts for the helicase genes. To note that *rhlE* helicase genes have two promoters, P1 and P2. The transcript from the P1 promoter is observed at low temperature (4°C), and that from the P2 promoter is produced at optimum temperature (22°C) of growth. The transcript start sites were based on transcriptome data of the laboratory (M. K. Ray, unpublished).

**Fig 2a.**
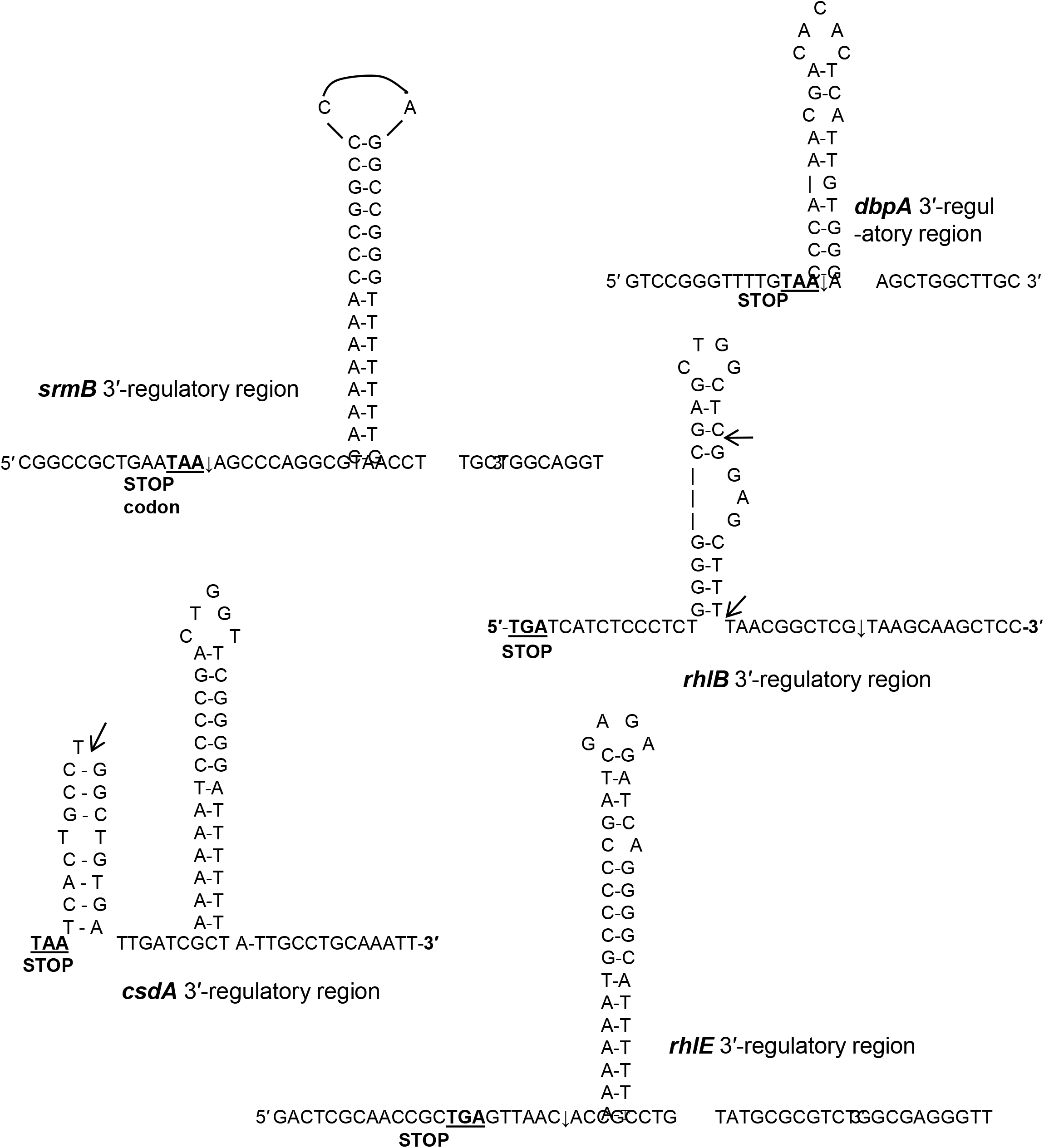
Regulatory 3’ regions of *P. syringae* helicase genes showing the putative 3’ end hairpin structures for transcription termination. The translational ‘Stop” codon of the genes are underlined. The arrows indicate the transcription stop site, that were determined in a separate study in the laboratory (M. K. Ray, unpublished data)

### Variation among *P. syringae* DEAD box RNA helicases and their similarities to the homologues from other *Pseudomonas* species

The five major DEAD box RNA helicases *rhlE*, *srmB*, *csdA*, *dbpA* and *rhlB* encode the proteins of different size, i.e., RhlE (618 residues), SrmB (448 residues), CsdA (557 residues), DbpA (461 residues) and RhlB (496 residues) corresponding to molecular mass of approximately 68 kDa, 49 kDa, 61 kDa, 50 kDa and 54 kDa respectively. They all displayed the conserved domain organization typical for the DEAD-box RNA helicases found in different organisms. (Fig 3a). Highly conserved catalytic ‘helicase core’ comprising the D1 and D2 domains, and variable N- and C-terminal extensions of the *P. syringae* RNA helicase proteins are shown schematically in Fig.3b.The N-terminal extensions are about 43 to 49 amino acids long except for the RhlB which showed an extension of ~140 amino acids. The C-terminal extensions were also of different lengths containing variable number of positively charged amino acid residues, which might have a bearing on interaction of the proteins with the negatively charged RNA substrates. The catalytic core of RNA helicase contains 12 conserved motifs, that are responsible for ATPase and helicase activities of the protein.

**Fig. 3.**
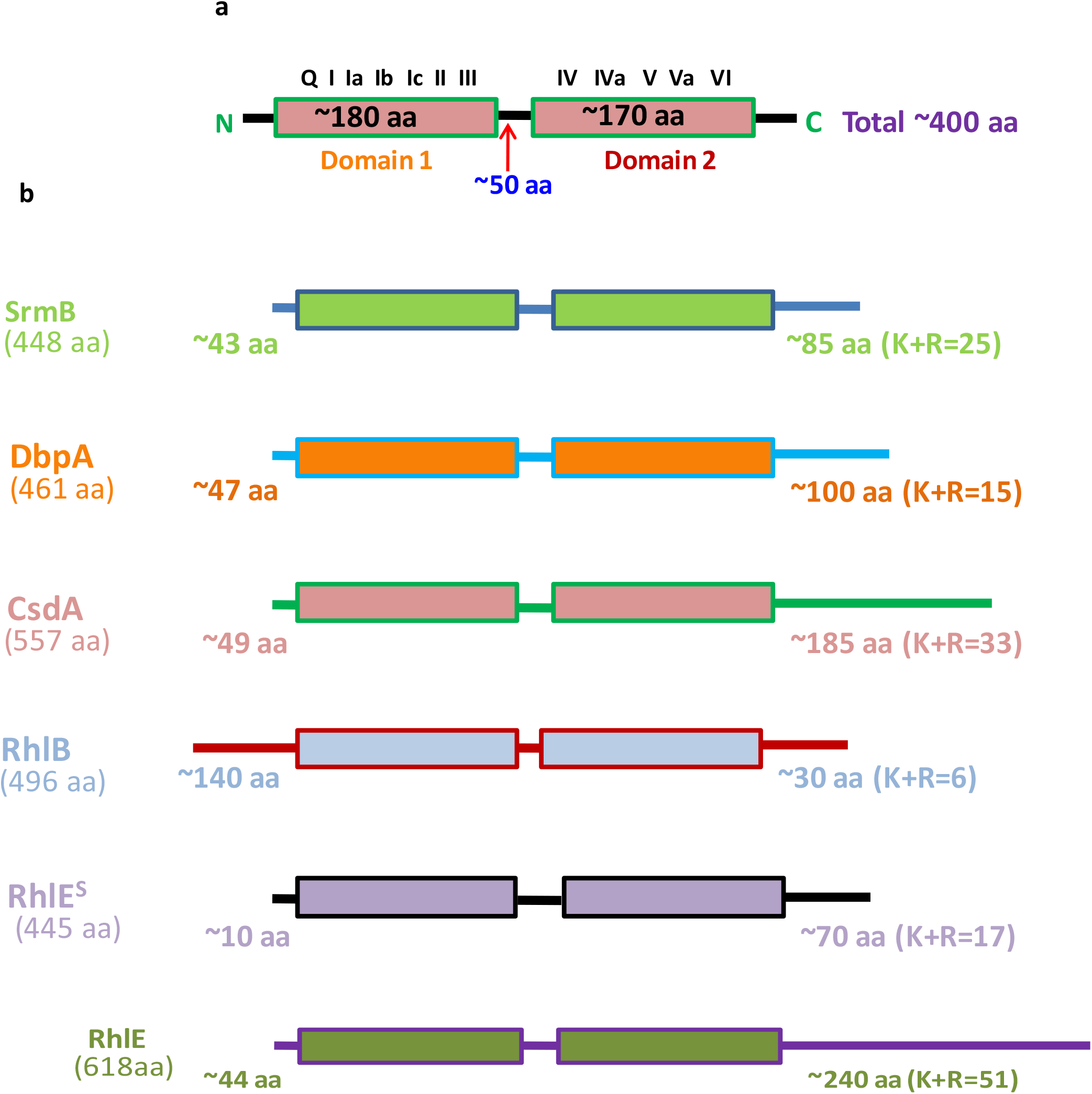
Domain organization of DEAD box RNA helicases: **(a)** Schematic representation of a typical RNA helicase showing twin domain catalytic center and linker. The domains are flanked by variable N and C terminal regions. Also shown in picture are different and highly conserved motifs involved in ATP hydrolysis and unwinding activities whereas the terminal regions are involved in binding substrate **(b)** Diagram representing number of amino acids, N and C terminal variable extensions and K+R of C-terminal extension in all six DEAD box RNA helicases of *P.syringae*.

Multiple sequence alignments of the six RNA helicase proteins of *P. syringae* Lz4W suggested that the helicases were about 32 to 39% identical among themselves in the amino acid sequences, with the lowest identity (32%) observed between CsdA and SrmB, and the highest (39%) shown by RhlB and RhlE (Fig 4 & Table 4a). The two RhlE homologues (RhlE-L) and RhlE-S), however had an amino acid identity of about 49.6% (Table 4a). An analysis with the *E. coli* RNA helicase homologues yielded similar results, except that the DbpA and CsdA helicases too, like the RhlB and RhlE, exhibited 39% identity. Thus, all the DEAD-box RNA helicases might have their common ancestry from which SrmB helicase might have diverged early in the evolution. In general, the respective *P. syringae* DEAD-box helicases show about 70 to 88 % identity with the *P. aeruginosa* homologues and 36 to 57% identity with the *E. coli* homologues in the amino acid sequences (Table. 4b). Thus, the *Pseudomonas* homologues of the RNA helicases have diverged substantially from the *E. coli* helicases while retaining similar cellular functions.

**Fig 4.**
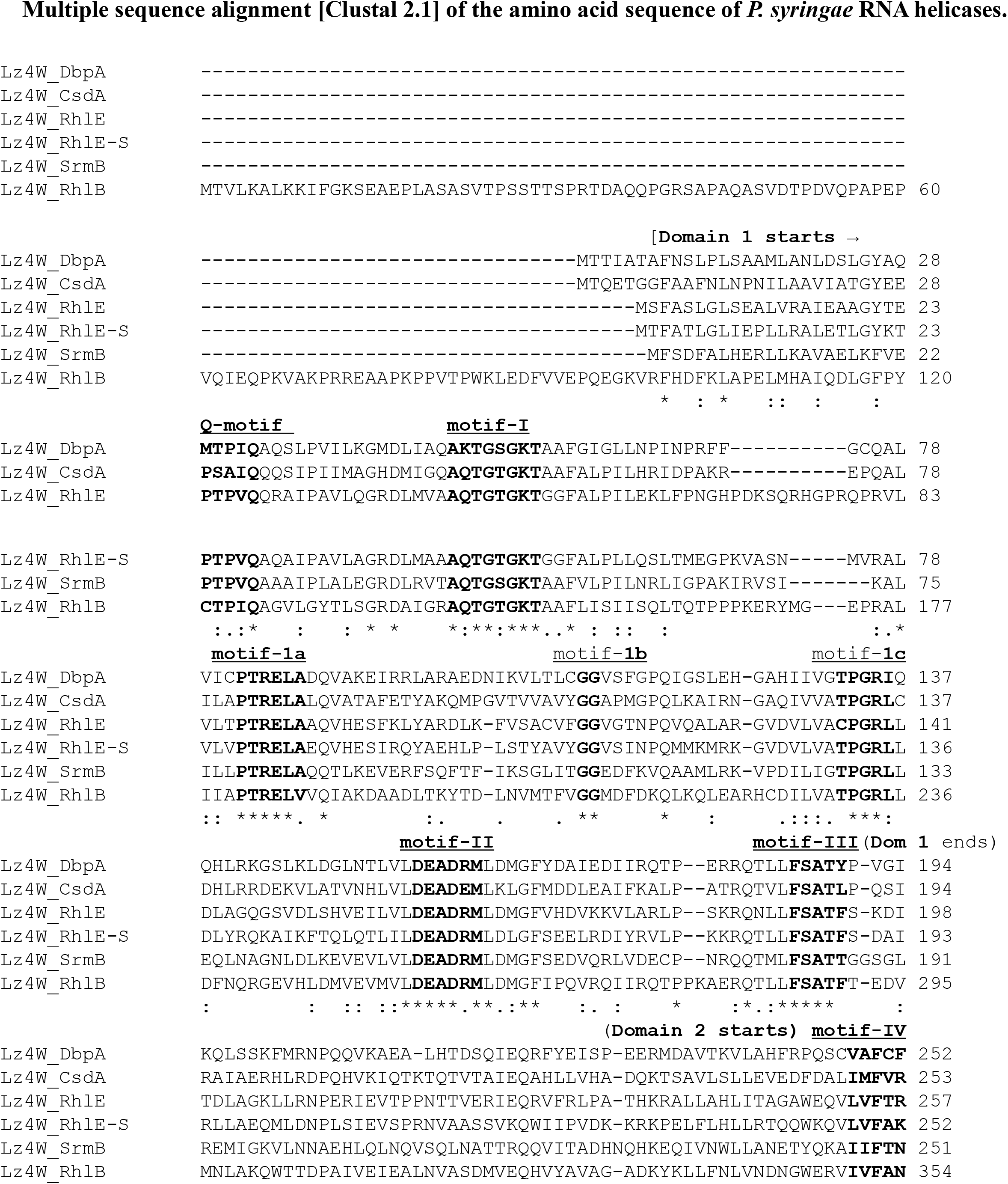
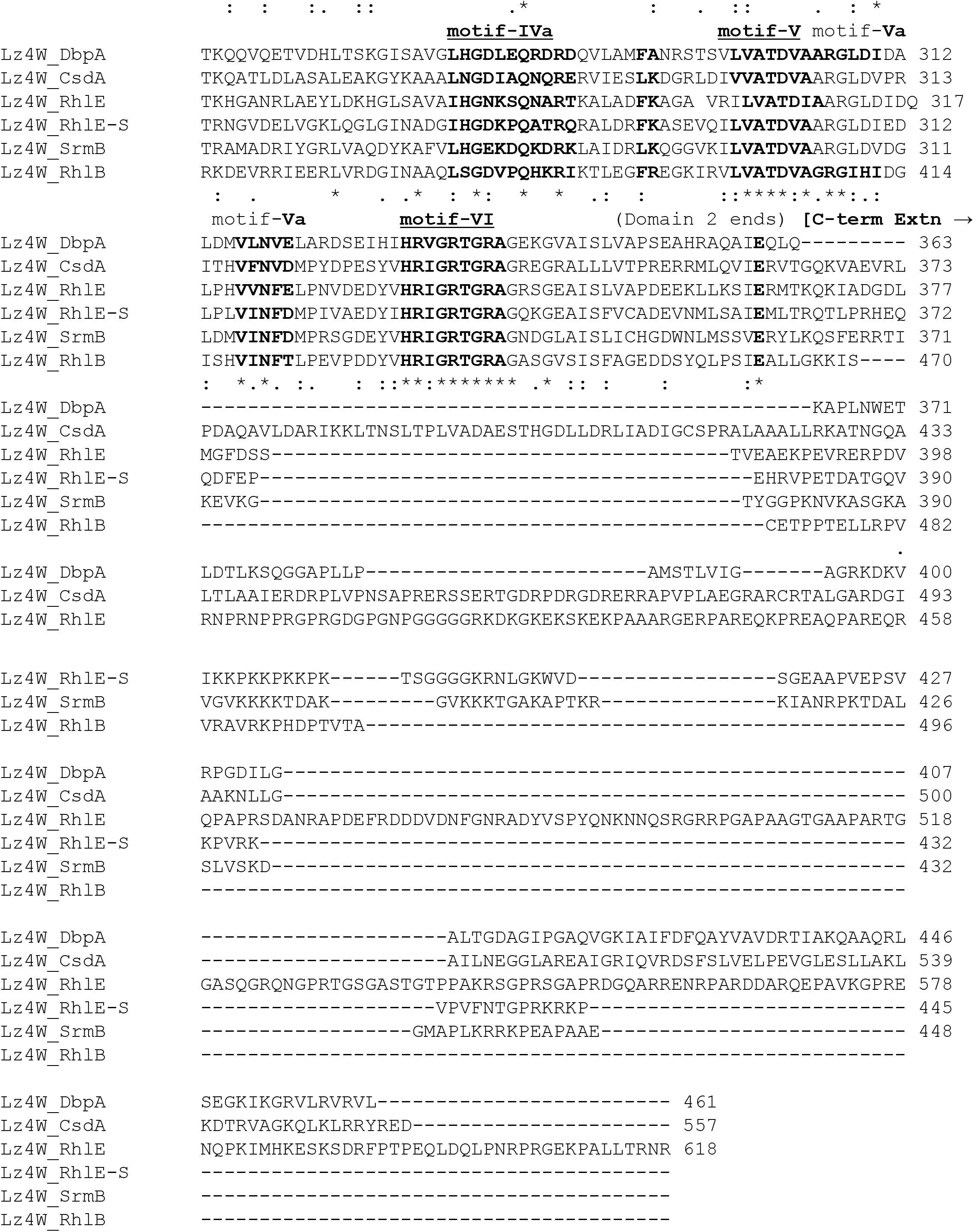
Multiple sequence alignment based analysis of DEAD box RNA helicases. Conserved structural motifs of the RNA helicases shown above the aligned amino acid sequences of the six RNA helicases encoded by the *P. syringae* Lz4W genome. The beginning and end of the two structural domains with C-terminal extensions of the RNA helicases have also been marked.

**Fig. 5.**
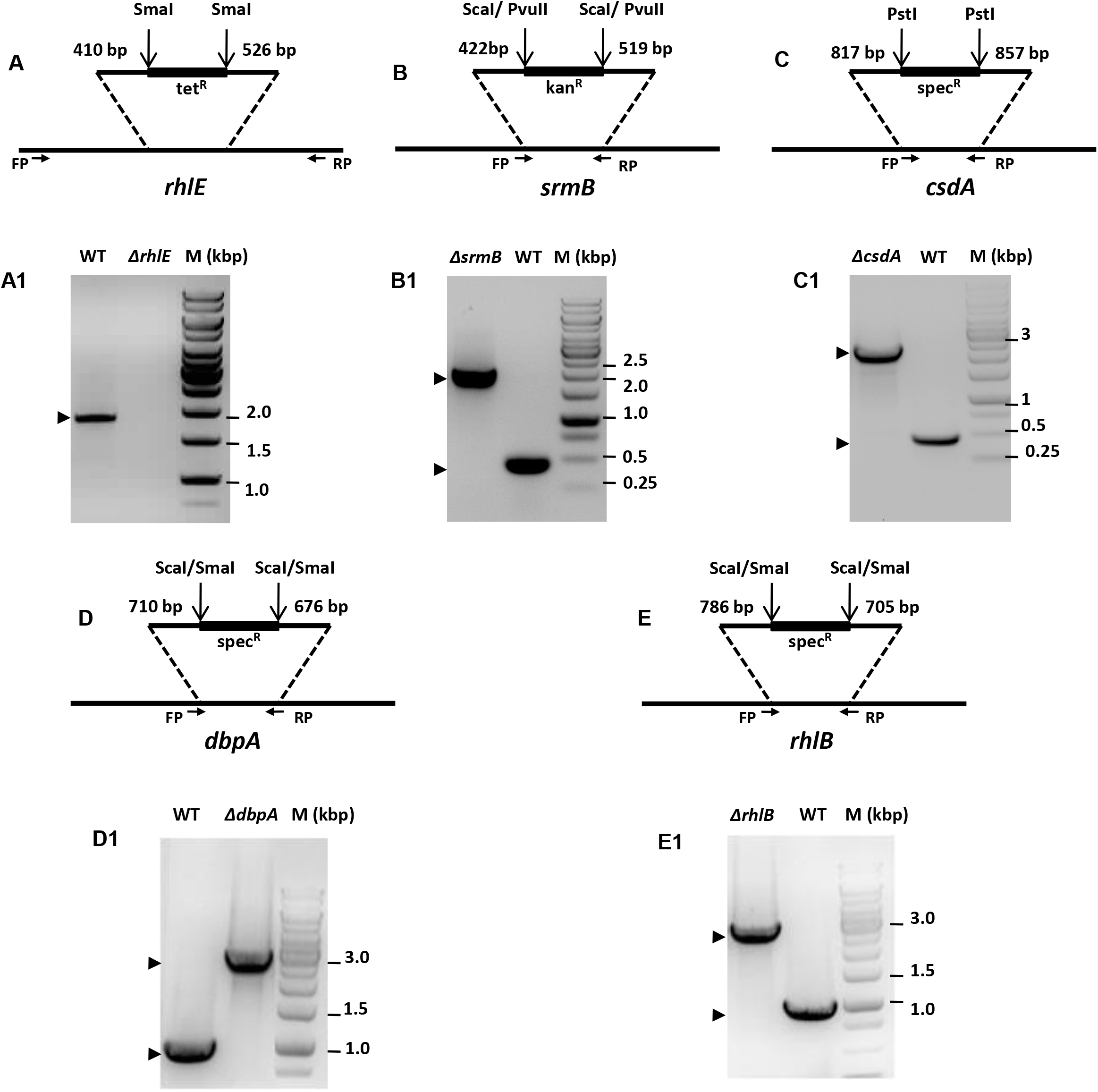
Generation and validation of RNA helicase mutants in *P. syringae*. Schematic representation of suicidal plasmid constructs employed for disruption of RNA helicase genes in wild type genetic background. Numbers above the gene in the schematic refer to nucleotide number of the plasmid born gene. Restriction sites used for insertion of selective marker are indicated. **(A)**Schematic representation of suicidal plasmid construct pJQ*rhlE-* tet used for disruption of *rhlE* gene. **(A1)**Lanes marked as WT and Δ*rhlE* represent the PCR amplification results of strain specific gnomic DNA with *rhlE* specific primers designed from each end of the *rhlE* gene. Reaction conditions for PCR were optimized to get amplification with wild type genomic DNA only.**(B)**Schematic representation of suicidal plasmid construct pJQ*srmB-* kan used for disruption of *srmB* gene. **(B1)**Lanes marked as WT, Δ*srmB* represent the PCR amplification results of strain specific gnomic DNA with *srmB* specific primers designed 225 base pairs from each end of inserted kan^r^ cassette. **(C**) Schematic representation of suicidal plasmid construct pJQ*csdA-* spec used for disruption of *csdA* gene. **(C1**) Lanes marked as WT, Δ*csdA* represent the PCR amplification results of strain specific gnomic DNA with *csdA* specific primers designed 250 base pairs from each end of spec^r^ cassette. **(D)**Schematic representation of suicidal plasmid construct *pJQdbpA-* spec used for disruption of *dbpA* gene. **(D1)**Lanes marked as WT, Δ*dbpA*, represent the PCR amplification results of strain specific gnomic DNA with *dbpA* specific primers designed 500 base pairs from each end of inserted spec^r^cassette.**(E)**Schematic representation of suicidal plasmid construct pJQ*rhlB-* spec used for deletion of *rhlB* gene. **(E1)**Lanes marked as WT and **Δ*rhlB*** represent the PCR amplification results of Strain specific gnomic DNA with *rhlB* specific primers designed 500 base pairs from each end of inserted spec^r^ cassette.

**Table 3.** Effects of helicase mutations on cell viability [in 4°C shifted cultures] and generation time of mutant strains grown at 22°C and 4°C. Dead cells were scored by counting the PI stained cells (red) and expressed as the % of total cells in the microscopic fields from minimum three different areas.

| Strains | Dead cells (%) at 4°C |  |  | Generation time (hours) |  |
| --- | --- | --- | --- | --- | --- |
|  | 24 hrs | 48 hrs | 72 hrs | at 22°C | at 4°C |
| WT | 0.2±0.11 | 1.5±0.08 | 1.4±0.2 | 2.02±0.21 | 6.75±0.5 |
| $\Delta csdA$ | 41.0±5.02 | 50.5±4.9 | 52.2±1.6 | 3.97±0.84 | *No growth |
| $\Delta srmB$ | 21.8±0.25 | 29.8±0.75 | 48.3±0.9 | 2.03±0.14 | 13.0±1.2 |
| $\Delta dbpA$ | 6.6±0.23 | 4.8±0.4 | 3.2±0.2 | 2.4±0.47 | 11.9±0.7 |
| $\Delta rhlB$ | 0.5±0.14 | 2.6±0.17 | 3.2±0.2 | 2.07±0.17 | 8.6±0.9 |
| $\Delta rhlE$ | 1.8±0.2 | 2.1±0.6 | 2.2±0.1 | 2.1±0.13 | 7.4±1.6 |

**Table 4.**
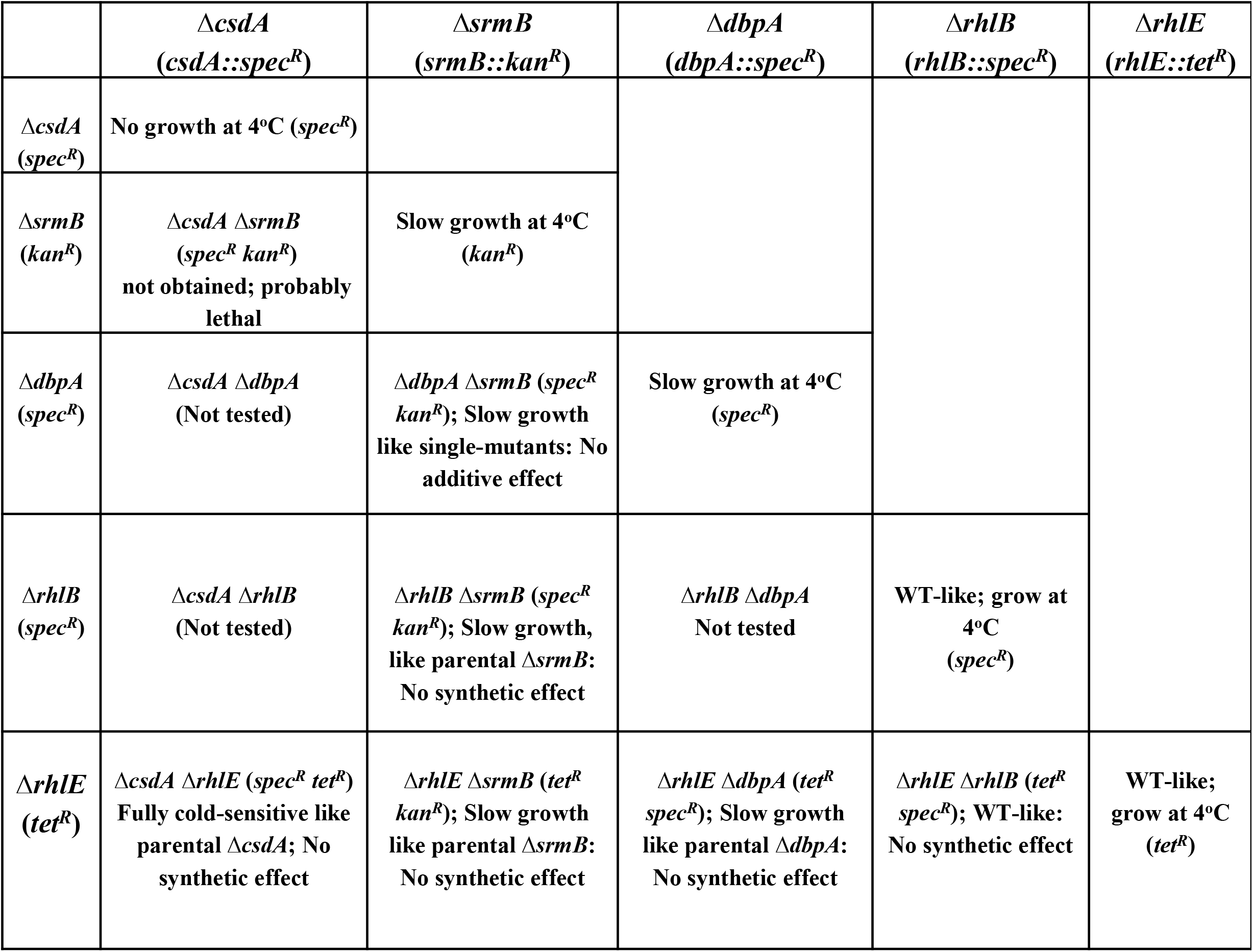
Genetic interaction among the helicase mutations of *P. syringae*, as judged from their combinational effects on the growth of the mutants at 4^°^C.

**Table 4a.** Identity among the six RNA helicases that are encoded by *P. syringae* Lz4W.

**# Percent Identity Matrix - created by Clustal 2.1**
|  | <u>DbpA</u> | <u>CsdA</u> | <u>RhlE</u> | <u>RhlE-S</u> | <u>SrmB</u> | <u>RhlB</u> |
| --- | --- | --- | --- | --- | --- | --- |
| 1: **Lz4W_DbpA** | **100.00** | 36.66 | 32.75 | 37.25 | 33.58 | 35.42 |
| 2: **Lz4W_CsdA** | 36.66 | **100.00** | 32.62 | 35.08 | 32.65 | 34.26 |
| 3: **Lz4W_RhlE** | 32.75 | 32.62 | **100.00** | 49.66 | 36.32 | 39.14 |
| 4: **Lz4W_RhlE-S** | 37.25 | 35.08 | 49.66 | **100.00** | 36.45 | 38.36 |
| 5: **Lz4W_SrmB** | 33.58 | 32.65 | 36.32 | 36.45 | **100.00** | 35.57 |
| 6: **Lz4W_RhlB** | 35.42 | 34.26 | 39.14 | 38.36 | 35.57 | **100.00** |

**Table 4b.**
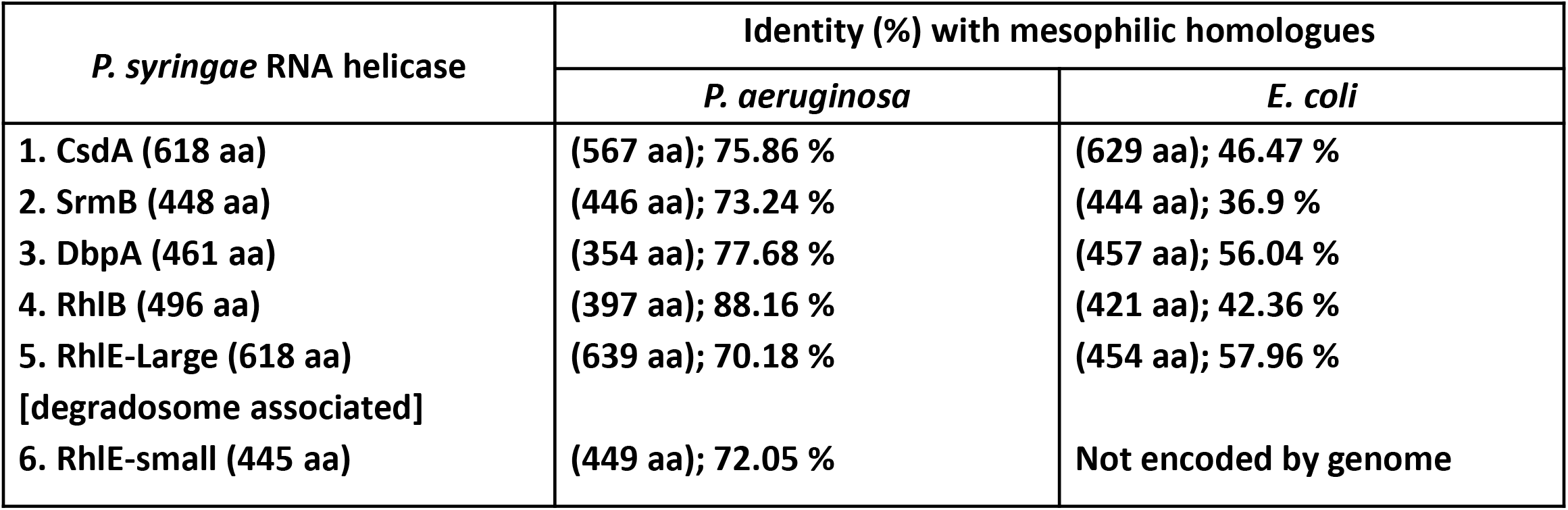
Percentage identity of DEAD box RNA helicases from the psychrophilic *P. syringae* Lz4W, and mesophilic *P. aeruginosa* and *Eschericha coli*.

### Importance of DEAD box RNA helicase gene deletions in cold adapted growth of *P. Syringae*

To decipher the role of the DEAD box RNA helicase genes in low temperature adapted growth of *P. Syringae*, mutant strains were constructed for each helicase gene by gene replacement through homologous recombination (Fig. 5, Table. 1). Growth analysis of mutant strains at 22°C revealed that Δ*rhlE*, Δ*srmB*, Δ*dbpA* and Δ*rhlB* strains displayed growth phenomenon similar to that of the WT, whereas growth of the Δ*csdA* strain was slightly impaired (Fig. 6). Measurement of generation times confirmed that Δ*rhlE*, Δ*srmB*, Δ*dbpA* and Δ*rhlB* have same growth rates as of wild type, however the generation time of Δ*csdA* (3.97 hrs) is almost twice that of wild type (2.02 hrs) (Table 3). However, growth study of all five mutants at low temperature (4°C) divulged interesting results. Helicase mutants Δ*rhlE* and Δ*rhlB* showed growth characteristics like wild type, however mutants Δ*srmB* and Δ*dbpA* displayed impaired growth, whereas Δ*csdA* mutant displaying a completely cold sensitive phenotype, did not exhibited any measurable growth (Fig. 7). Measurement of generation times confirmed that Δ*rhlE*, Δ*rhlB* have same growth rate as of wild type, however the generation time of Δ*srmB* (13.0 hrs) and Δ*dbpA* (11.9 hrs) is significantly longer than that of wild type strain (6.75 hrs). The results of growth analysis for *P. Syringae* strains Δ*csdA* (22°C and 4°C) and Δ*dbpA* (4°C) were different to results obtained with mesophilic *E. Coli*, where Δ*csdA* only exhibits slow growing phenotype at low temperature (25°C). These findings indicate that helicase genes *srmB* and *dbpA* are important but *csdA* is indispensable for growth of bacterium at low temperature (4°C).

**Fig. 6.**
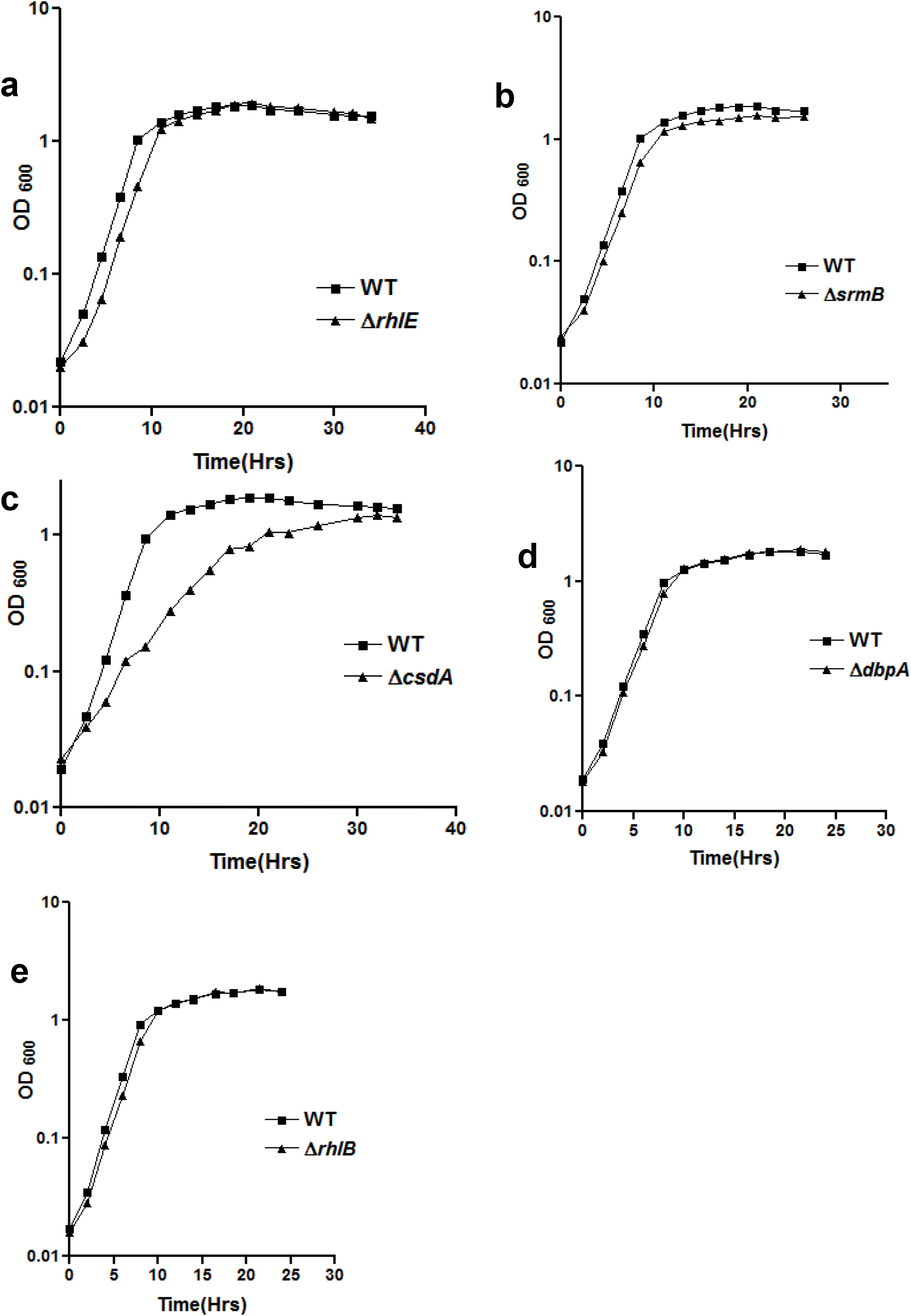
Growth analysis of *P. syringae* helicase mutants Δ*rhlE* (a), Δ*srmB* (b), Δ*csdA* (c), Δ*dbpA* (d) and Δ*rhlB* (e) at 22°C. For measurement of growth, mutant strains of *P. syringae* were grown separately in ABM broth at 22°C and OD at 600nm [OD_600_] was recorded at regular time intervals and plotted against time. All growth curves were generated using GraphPad Prism 4.0 software.

**Fig. 7.**
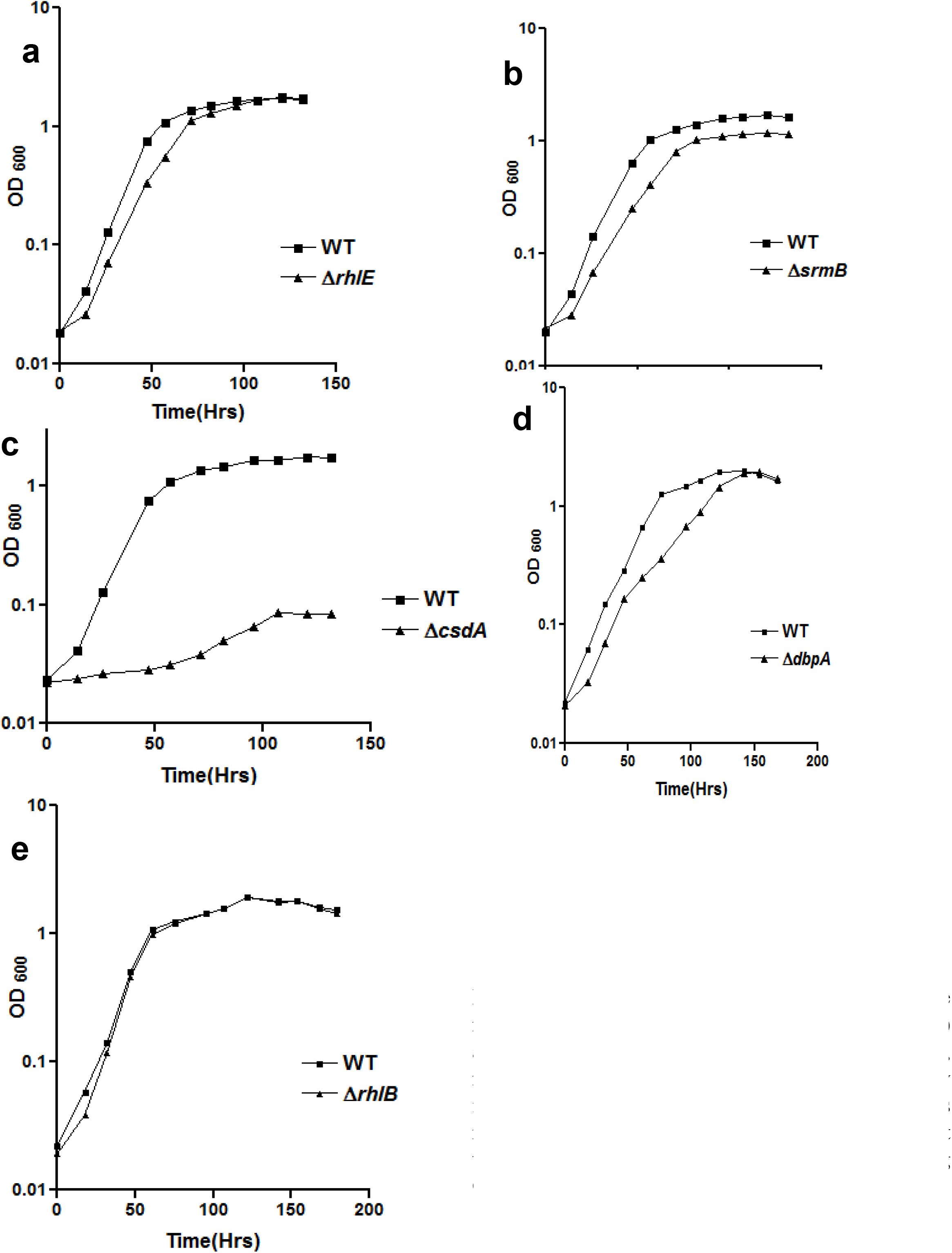
Growth analysis of *P. syringae* helicase mutants Δ*rhlE* (a), Δ*srmB* (b), Δ*csdA* (c), Δ*dbpA* (d) and Δ*rhlB* (e) at 4°C. For measurement of growth, mutant strains of *P. syringae* were grown separately in ABM broth at 4°C and OD at 600nm [OD_600_] was recorded at regular time intervals and plotted against time. All growth curves were generated using GraphPad Prism 4.0 software.

**Fig 7.**
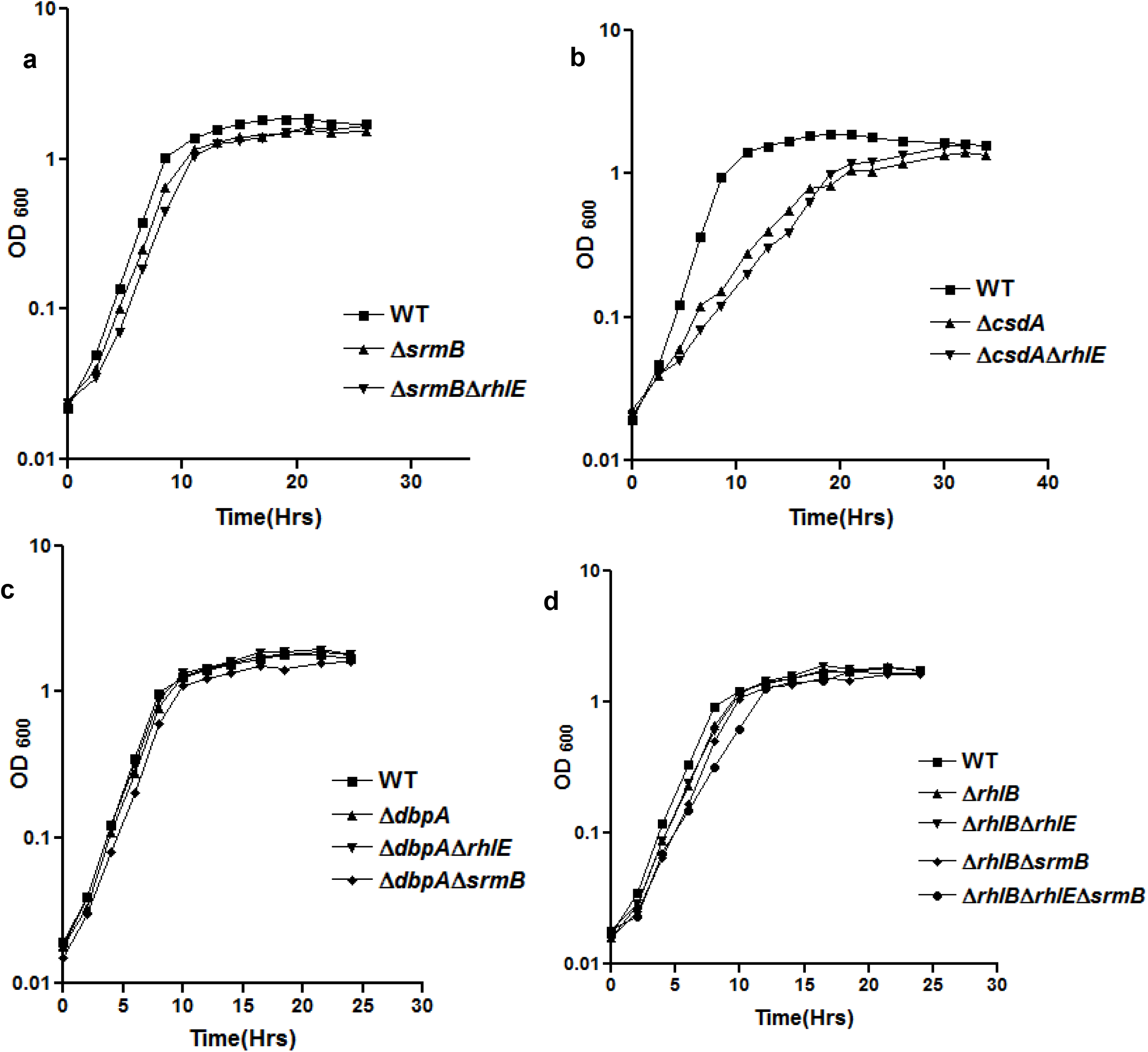
Growth analysis of DEAD box RNA helicase mutants harbouring multiple disruptions. Growth analysis of *srmB* (**a**) and *csdA* (**b**) gene disruptions in different mutant backgrounds was performed at 22°C. Growth of Δ*dbpA*, (Δ*dbpA*Δ*rhlE*) and (Δ*dbpA*Δ*srmB*) strains at 22°C are shown in (**c**). Similarly effects of *rhlB* deletion in various helicase mutant backgrounds was analyzed by comparing the growth profiles of Δ*rhlB*, (Δ*rhlB* Δ*rhlE*), (Δ*rhlB* Δ*srmB*), and (Δ*rhlB* Δ*rhlE* Δ*srmB*) strains at 22°C **(d).** For measurement of growth, different strains of *P. syringae* were grown separately in ABM broth at 22°C. OD at 600nm [600_nm_] was recorded at regular intervals and plotted against time. All growth curves were generated using GraphPad Prism 4.0 software.

The growth phenotypes of RNA helicase mutant strains grown at 22°C or 4°C (After shift from 22°C) was further investigated by cell viability assays using Live-Dead staining of the cells in growing cultures. After 72 hours of incubation at 4°C, approximately 52% of Δ*csdA* cells and 48% of Δ*srmB* cells stained red (Dead), as compared to wild type where only 1% of cells stained red (Table. 1). The results obtained with Live-Dead staining of wild type and mutant strains is well in agreement with the results obtained from growth curve analysis of desired mutant strains. The mutant strains with increased generation time at low temperatures displayed high percentage of cell death at low temperatures. The anomaly was with Δ*dbpA* strain, that displayed a prolonged generation time (11.9 hours) similar to Δ*srmB* strain (12.9 hours), however the percentage of dead cells was very less than expected. The low percentage of cell death may be due to better acclimatization of Δ*dbpA* to low temperature (4°C) after shift from optimal conditions of growth (22°C).

Microscopic studies of helicase disrupted strains revealed a decrease in the cell size as one of the major effects of exposure to low temperature (4°C), which has also been reported earlier [25, 26]. Like WT cells, Δ*rhlE* and Δ*rhlB* strains did not display any low temperature associated change in morphology (Fig S2). Δ*csdA* mutants grown at 22°C displayed a dull appearance whereas Δ*csdA* cells grown at 4°C displayed a speck of dark materials at the middle of the lateral along long axis, giving an appearance of puncture at center of the cells [Fig S2]. Helicase mutants Δ*srmB* and Δ*dbpA* and did not exhibit any changes in appearance at 22°C, however when these mutants were grown at 4°C the mutants revealed a dull appearance and were fractionally elongated and bigger in size than wild type cells growing at same (4°C) temperature. The low temperature associated morphological changes observed with Δ*srmB*, Δ*csdA* and Δ*dbpA* mutants might be an indirect effect of *srmB, csdA* or *dbpA* gene disruption in mutant cells or these genes might have an indirect role in modulation of cell shape and size in *P. syringae* especially at low temperature. The magnitude of low temperature associated changes in morphology with Δ*srmB* and Δ*dbpA* mutants was less pronounced as compared to effects seen with Δ*csdA*.

### Growth phenotypes of the double and triple helicase deletion strains

The disruption of *rhlE* gene in Δ*srmB* background generating the double mutant (Δ*rhlEΔsrmB*)did not lead to any rescue of the slow-growing phenotype in Δ*srmB* mutant of *P. syringae*, as observed in the case of *E. coli* [12]. There was no difference in the growth profiles between Δ*srmB* and Δ*srmBΔrhlE* at low temperature (Fig. 8a). This suggested that the *P. syringae* RhlE helicase does not probably interact with the SrmB-depleted 50S ribosomal subunit, as observed in *E. coli* where RhlE was proposed to modulate the inter conversion of two alternative forms of 50S ribosomal subunit intermediates. SrmB requirement (possibly in the ribosome assembly) during the growth of psychrophilic *P. syringae* at low temperature is independent of RhlE function, mainly mediated through RNA degradosome in the bacterium. Similarly, but in difference to *E. coli*, both Δ*csdA* and Δ*csdAΔrhlE* double mutant of *P. syringae* displayed no alteration in the cold sensitive phenotype of Δ*csdA* mutant (Fig. 8b). In *E. coli*, deletion of *csdA* gene in Δ*rhlE* background exacerbated the cold sensitive phenotype of Δ*csdA*.

**Fig 8.**
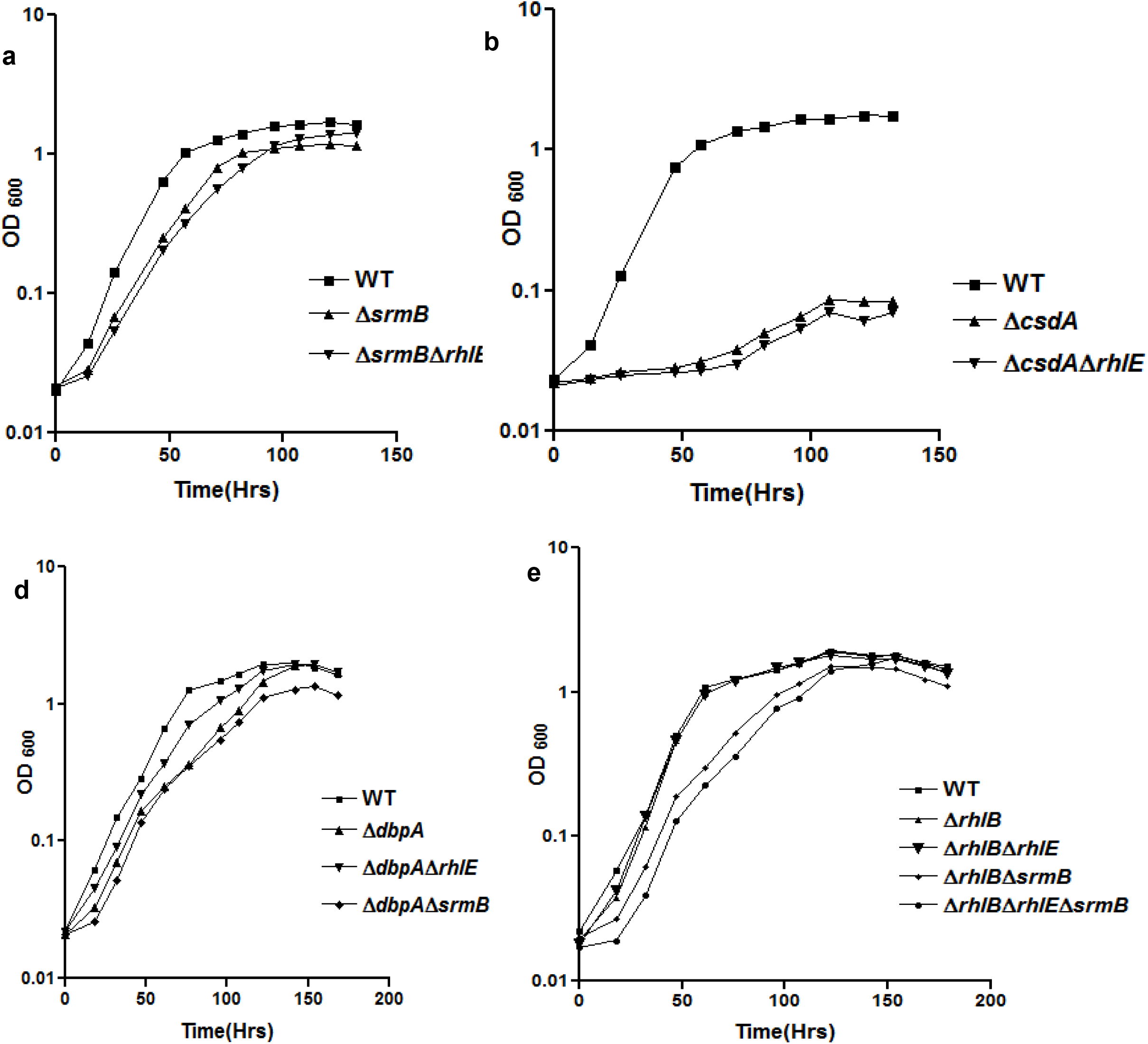
Growth analysis of DEAD box RNA helicase mutants harbouring multiple disruptions. Growth analysis of *srmB* (**a)**and *csdA* (**b**) gene disruptions in different mutant backgrounds was performed at 4°C. Growth study of Δ*dbpA, (ΔdbpAΔrhlE*) and (Δ*dbpAΔsrmB*) strains at 4°C is shown in (**c**). Similarly effects of *rhlB* deletion in various helicase mutant backgrounds was analyzed by comparing the growth profiles of Δ*rhlB, (ΔrhlB ΔrhlE), (ΔrhlB ΔsrmB*), and (Δ*rhlB ΔrhlE ΔsrmB*) strains at 4°C (**d**). For measurement of growth, different strains of *P. syringae* were grown separately in ABM broth at 4°C. OD at 600nm [600_nm_] was recorded at regular intervals and plotted against time. All growth curves were generated using GraphPad Prism 4.0 software.

Disruption of *dbpA* gene in wild-type, Δ*rhlE* and Δ*srmB* genetic backgrounds has some interesting effects on the growth and viability of the strains. Growth of Δ*dbpA, ΔdbpAΔrhlE* and Δ*dbpAΔsrmB* strains at 22°C and 4°C are shown in (Fig.7c and 8c). All strains including the wild-type, Δ*dbpA*, Δ*dbpA*Δ*rhlE* and Δ*dbpA*Δ*srmB* displayed normal and similar growth at 22°C (Fig. 7c). However, at 4°C, while Δ*dbpA* displayed a marginally cold-sensitive phenotype, the *dbpA* deletion in Δ*rhlE* background ameliorated the cold-sensitive phenotype of Δ*dbpA* strain. The Δ*dbpAΔrhlE* double mutant grew better than the single Δ*dbpA* mutant (Fig. 8c). On the other hand, the double mutant Δ*dbpA*Δ*srmB* displayed cold sensitive phenotype similar to the single mutants Δ*dbpA* and Δ*srmB*. This indicates that, although both genes are important for growth at low temperature, there is no additive effect on the severity of cold-sensitive phenotype in the double mutant Δ*dbpA*Δ*srmB* when both helicases are missing from the cells.

The effects of *rhlB* deletion in various helicase mutant backgrounds was analyzed by comparing the growth profiles of Δ*rhlB, ΔrhlB ΔrhlE, ΔrhlBΔsrmB*, and Δ*rhlB ΔrhlEΔsrmB* strains at 22°C and 4°C (Fig. 7d and 8d). All these strains displayed normal growth at 22°C, and no appreciable differences were observed in their growth profiles (Fig. 7d). At 4°C, Δ*rhlB* and Δ*rhlBΔrhlE* strains displayed wild type like growth, suggesting that the two mutations exerted no additive effects, and both of them are dispensable for growth at low temperature singly or simultaneously. It is also important to note that the double mutant Δ*rhlBΔsrmB* grew slowly at 4°C, similar to the cold sensitive single Δ*srmB* mutant, suggesting that RhlB depletion did not exacerbate the growth defect of SrmB depleted cells (Fig 8d). This was akin to the results obtained with RhlE depleted cells of Δ*srmB* in Δ*rhlEΔsrmB* double mutant (Fig. 8a). A summary of the interactions between the various helicase deletions of *P. syringae* are presented in Table 4.

Remarkably, severity of the cold sensitive growth defect in the triple mutant Δ*rhlBΔrhlEΔsrmB* (Fig. 8d) lacking RhlB, RhlE, and SrmB helicases simultaneously, was enhanced compared to the double mutants Δ*rhlBΔsrmB* (Fig. 8d) or Δ*rhlEΔsrmB* (Fig. 8b).The modest but marginal increase in the severity of the defect in the triple mutant at low temperature suggested that, although RhlB and RhlE helicases are ordinarily dispensable for the growth, they play certain subtle role in RNA metabolism whose effects are manifested during the growth only when both of them are removed together with SrmB in the cells at low temperature.

### Functional complementation of RNA helicase mutant strains

Genetic complementation of the helicase deletion mutants was important for confirmation that the growth defects in the mutants are only due to gene disruption and not due to any other unexpected secondary alteration in the cells. This would also confirm the theoretical prediction based on genome analysis, that there will be no polar effects on downstream genes in the mutants, due to disruption of the monocistronic helicase genes. Additionally, cross complementation of one helicase mutants with other helicases will provide an insight into the functional redundancy between the DEAD box RNA helicases. Accordingly, each of the five helicase genes were cloned in the broad host range plasmid, pGL10, which can replicate in *P. syringae* Lz4W and the helicase genes are constitutively expressed from the *lacZ* promoter of the pGL10.

The complemented mutants were grown at 22°C and 4°C, for analyzing their growth properties compared to the parental mutant strain. Since the copy number of the pGL10 based plasmid is ~5 copies per cell in *P. syringae* (Laboratory observation), the level of expressed proteins were never more than 5 folds in the cells (data not shown). The 22°C and 4°C growth profiles of the complemented cold-sensitive strains Δ*csdA* (Fig. 9d, 10d), Δ*srmB* (Fig. 9c & 10c), and Δ*dbpA* (Fig. 9e & 10e) show that the growth defects of the helicase mutants are rescued only by the cognate helicases expressed from the plasmids in the cell. The cold sensitive phenotype of helicase mutants could not be rescued by expressing others helicases, suggesting that there is no cross complementation, or any partial rescue of the phenotypes due to overlapping functions.

**Fig. 9.**
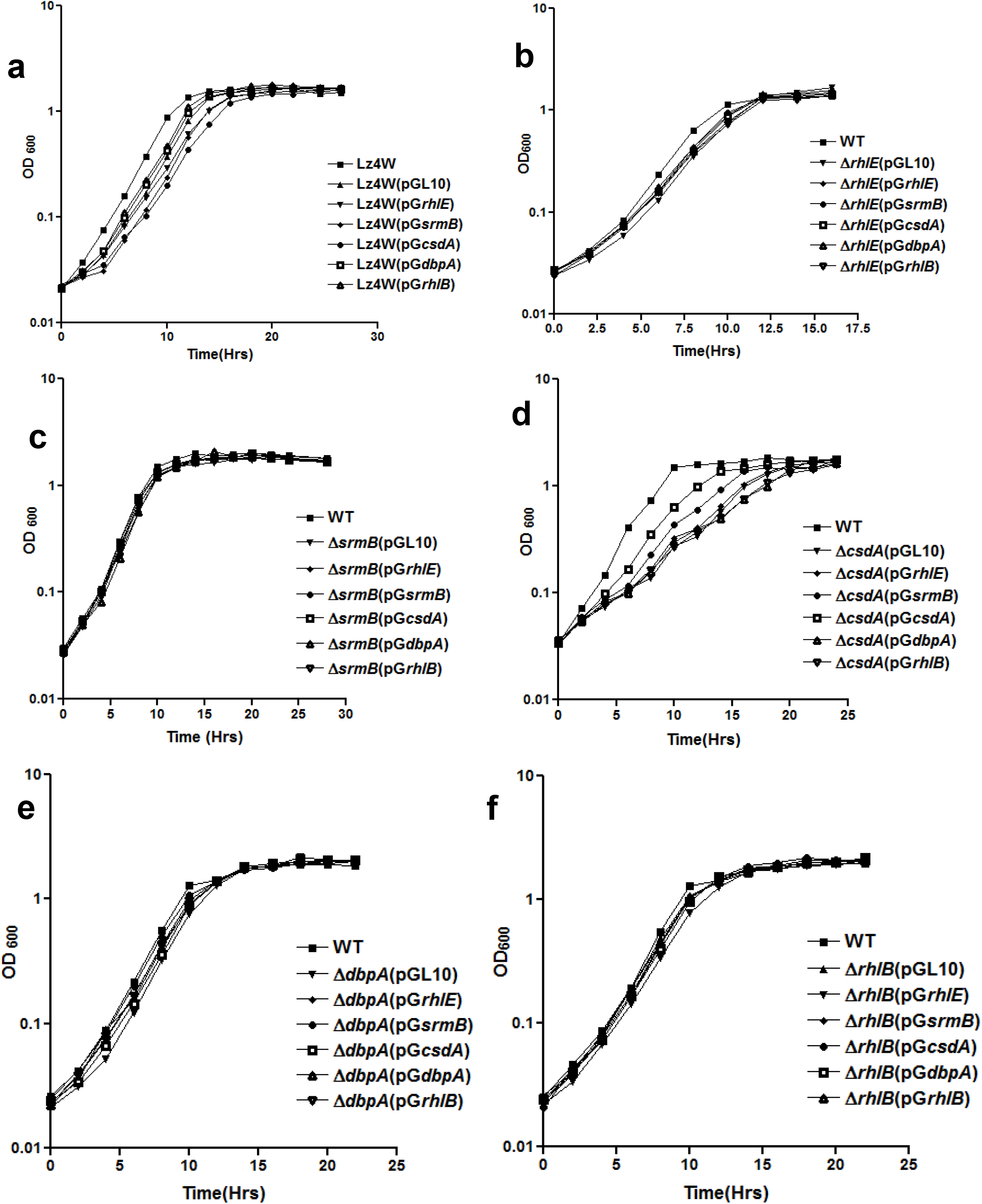
Functional complementation of helicase mutant strains: Growth analysis for overexpression of all five plasmid born helicases in WT [Lz4W], Δ*rhlE, ΔsrmB, ΔcsdA, ΔdbpA and ΔrhlB* mutant strains at 22°C.

**Fig. 10.**
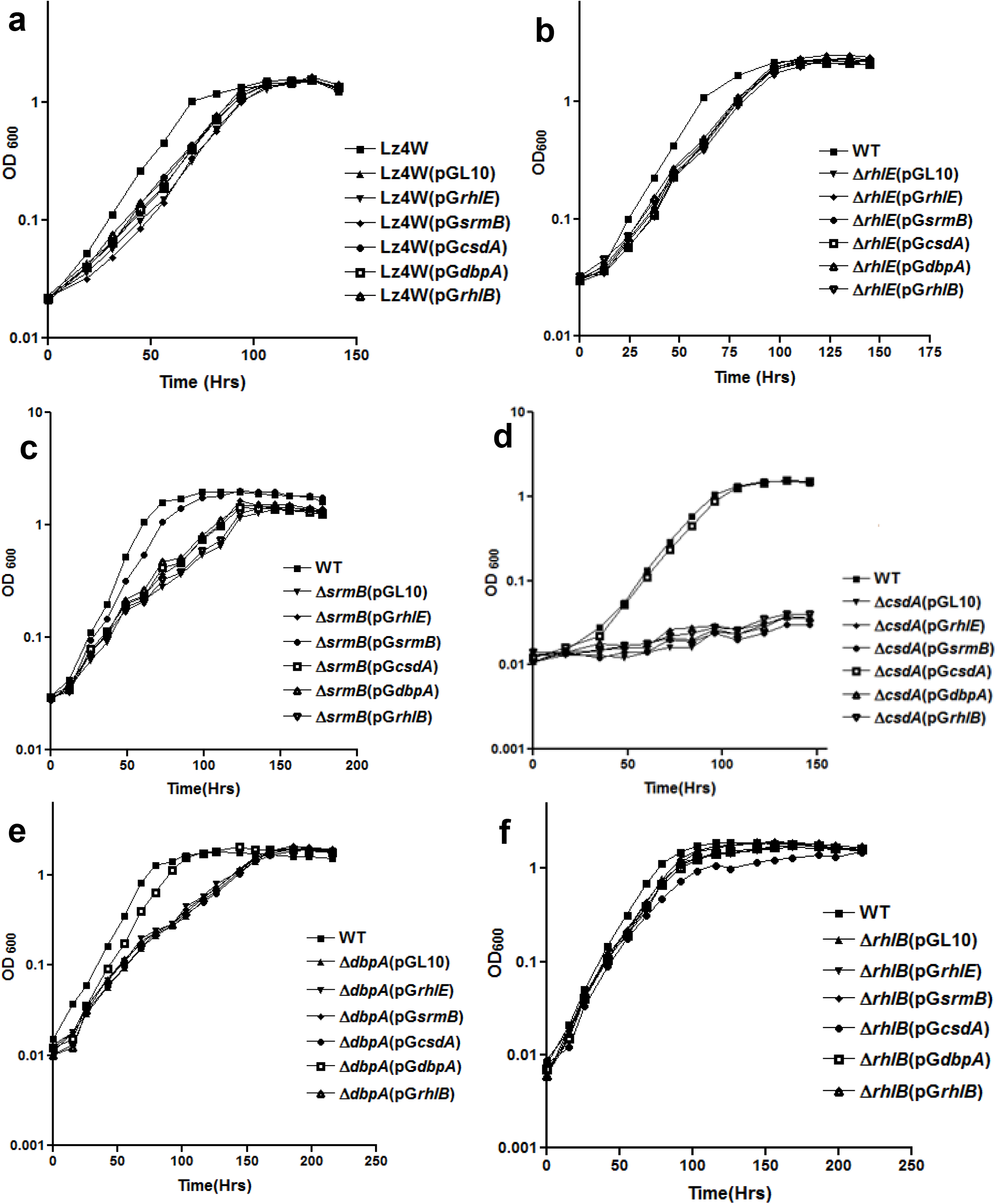
Functional complementation of helicase mutant strains: Growth analysis for over expression of all five plasmid born helicases in WT [Lz4W], Δ*rhlE, ΔsrmB, ΔcsdA, ΔdbpA and ΔrhlB* mutant strains at 4°C.

The complemented mutants Δ*rhlB* (Fig. 9f & 10f) and Δ*rhlE* (Fig. 9b & 10b) harbouring all six plasmids including the empty pGL10 also did not show any alteration in their growth due to over expression of the RNA helicases from the plasmids. Δ*rhlB* and Δ*rhlE* mutants by themselves also did not display any growth defects at either temperature (22°C and 4°C). Growth analysis of wild type strain harbouring the additional RNA helicase genes on plasmids did not reveal any effect either on the growth or on the viability of the cells at 22°C or 4°C (Figs. 9a & 10a).

## Discussion

Genomic analysis shows that the RNA helicase genes are not clustered in a specific chromosomal segment, but dispersed all over the bacterial chromosome. The helicase genes are monocistronic and regulated by their independent promoters and transcription termination signals. The nucleotide sequence of the regulatory region of these genes did not throw up any noticeable sequence motifs that are specific for these genes or their expression at low temperature. Only the *csdA* transcript was observed to have a long 5’-UTR (227 base pairs) similar to the low temperature specific transcript of *rhlE* that had 213 base pairs long 5’-UTR. But the two genes did not have any common regulatory sequences which can be correlated to their expression or their significance in the cells. Nonetheless, the monocistronic gene organization of helicase genes helped us to create gene knockout mutants by employing the antibiotic resistance cassette insertion without affecting the downstream gene expression (polar effects).

Analysis of the single deletion mutants of helicase genes has shed new light on the role of the individual RNA helicases in the growth of *P. syringae*. Our study shows that three of the five DEAD box RNA helicases are important for growth of the psychrophilic bacterium at low temperature (4°C). These three helicases are SrmB, CsdA and DbpA all of which are known to affect the biogenesis of 50S ribosomal subunit and hence mature 70S ribosomes in *E. coli* [15, 16]. Interestingly, the *srmB* and *csdA* deletions in mesophilic *E. coli* also led to a cold sensitive phenotype in the bacterium, in which Δ*srmB* and Δ*csdA* mutants could grow at 25°C (low temperature for the mesophile) with a longer generation time (~90 and 138 min, respectively), compared to the wild-type (generation time 77 min [12]. The Δ*dbpA* did not show any cold sensitive growth defect at 25°C. These are in contrast to our observation with the psychrophilic *P. syringae*, in which *csdA* is absolutely essential, and the *srmB* or *dbpA* genes are important for the growth at low temperature (4°C). The Δ*csdA* mutant did not grow at all, while Δ*srmB* and Δ*dbpA* mutants grew slowly at the low temperature. In this respect, the requirement of *dbpA*, and the absolute necessity of *csdA* for growth of the psychrophile at low temperature (4°C) are novel findings of this study.

At optimal temperature for growth, the requirement of DEAD box RNA helicases has not been reflected by mutational studies in bacteria, especially using the single deletion mutants. At 37°C, all single helicase-deletion mutants of *E. coli* exhibited ~25.5 minutes generation time, similar to wild-type [12]. Interestingly, Δ*csdA* mutant of *P. syringae* demonstrated the importance of *csdA* at the optimal temperature (22°C) of growth for the psychrophilic bacterium. The Δ*csdA* showed a substantial increase in the generation time (3.97 hrs) as compared to wild type (generation time, 2.02 hrs). All other four single deletion mutants for DEAD box RNA helicases (Δ*rhlE*, Δ*srmB*, Δ*dbpA* and Δ*rhlB*) did not show any growth defect or alteration in the viability at optimum temperature (22°C). These results suggested that the various RNA folding pathways and the RNA secondary structures are optimized for functions at the temperature in which bacteria grows at a maximum rate. The lack of any particular RNA helicases is not probably felt unless the RNA helicase is necessary for a very specific function(s) that affects cell viability or cell growth. The *csdA* perhaps does similar function in *P. syringae*, as a result of which Δ*csdA* mutant grows slowly at 22°C and fails to grow due to cellular lethality at lower temperature (4°C). [42]

The loss of cell viability at low temperature due to RNA helicase deficiency was observed not only with Δ*csdA* but also in two other cold sensitive mutants (Δ*srmB* and Δ*dbpA*) of *P. syringae*. These mutants grew slowly (generation time 13.01 hours for Δ*srmB* and 12.6 hours for Δ*dbpA}* compared to the wild-type (generation time 6.75 hours) at 4°C, suggesting that the slow growing phenotype could be partly related to the cell death observed in the mutants at low temperature. The slower RNA metabolism in the absence of RNA helicases is likely to affect the growth rate, mostly at the low temperature. Role of *csdA* and *srmB* in low temperature growth of mesophilic *E. Coli* has been reported earlier, however the importance of *dbpA* in cold adapted growth of *P. Syringae* was unique finding of the study.

Due to functional redundancy of gene products, especially in multigene families, the mutant phenotype of the cells is not manifested sometimes in the single deletion mutants. The double deletion mutants were constructed using the disrupted alleles of Δ*rhlE, ΔsrmB, ΔcsdA, ΔdbpA* and Δ*rhlBΔcsdA*, which indicated that there is not much additive or synergistic effects due to their combination in the cells, except for the double mutants Δ*srmBΔcsdA* and Δ*dbpAΔrhlE*. The Δ*srmBΔcsdA* mutant could not be recovered in our experiments, possibly due to the combinatorial lethality. On the other hand, the double mutant Δ*dbpAΔrhlE* grew marginally better than the slow growing Δ*dbpA* at the low temperature (4°C), suggesting a possible interaction between RhlE and DbpA on an unidentified common substrate (RNA) in the cells, in which RhlE exacerbates the effects of DbpA depletion (in Δ*dbpA}*. When RhlE is removed from such cells, as in Δ*dbpAΔrhlE*, the deleterious effect of DbpA depletion is marginally relieved in the double mutant. We also observed that, although Δ*rhlBΔrhlE* does not have a growth defect, and Δ*rhlBΔsrmB* and Δ*rhlEΔsrmB* combinations do not show ameliorating (synthetic rescue) or worsening (synthetic lethality) of the cold-sensitive phenotype of Δ*srmB*, the triple-deletion Δ*rhlBΔrhlEΔsrmB* exacerbates the cold sensitive growth of Δ*srmB*. These results suggest that the cellular lack of RhlB or RhlE activities are not manifested in the cell growth, possibly due to their subtle roles in cellular milieus of Δ*srmB*, but when all of them are combined in Δ*srmB* background, their requirements are manifested in the growth rate of the mutant, which slows down further at the low temperature. The results of these mutational studies thus point towards the existence of different types of subtle and not-so-subtle interactions among the RNA helicases in *P. syringae*.

Microscopic studies of the DEAD-box RNA helicase depleted cells suggested a role for the RNA helicases in regulation of cell size and morphology, as evidenced by the alteration in cell morphologies of the cold-sensitive Δ*srmB*, Δ*csdA* and Δ*dbpA* mutants at low temperature. However, these effects are unlikely to be direct by a lack of the helicase proteins, and possibly resulted from the affected pathways that determine cell morphology or cell size.

Our results suggest that the DEAD box RNA helicase *csdA* is absolutely necessary for the psychrophilic adaptation of the Antarctic *P. Syringae*. The study also pointed out the importance of *dbpA* helicase in the cold-adaptation of the psychrophilic bacterium, which binds to a specific region (helix 92) of 23S rRNA but the deletion of which does not affect the growth of mesophilic *E. coli* at low temperature [17, The current study also pointed out the importance of *srmB* helicase at low temperature, which has been implicated in the assembly of 50S ribosomal particle in *E. coli* [15, 16]. Thus, all the three DEAD box RNA helicases which have been found to be important for growth of the Antarctic *P. Syringae* at low temperature are known to participate in the 50S ribosomal subunit maturation and ribosome biogenesis in *E. coli* and *B. Subtilis* [11, 14–17], suggesting the crucial dependence of ribosome assembly and protein synthesis on RNA helicases at low temperature.

## Acknowledgments

The authors acknowledge Council for Scientific and Industrial Research (CSIR), Government of India and Indian Council for Medical Research (ICMR), Government of India for financial support in the form of fellowship to Ashaq Hussain during this study.

## Author contributions

Conception and design of study: Ashaq Hussain and Malay K Ray

Resources: Malay K. Ray

Supervision: Malay K. Ray

Methodology: Ashaq Hussain

Data collection and interpretation: Ashaq Hussain

Drafting of article: Ashaq Hussain

Review & editing: Ashaq Hussain

## Conflict of interest

There are no conflicts to declare.

## Supplementary information to

**Table S1.** Plasmids for high level expression of RNA helicases in BL21 cells.

| Plasmid | Description | Reference/Source |
| --- | --- | --- |
| pET28a | *Kan<sup>r</sup>*, Expression vector for N-terminal His-tagged proteins | Novagen |
| pET28arhlE | pET28a plasmid for over-expression of N-terminal His tagged RHLE of *P. syringae*. | This study |
| pET28asrmB | pET28a plasmid for over-expression of N-terminal His tagged SRMB of *P. syringae*. | This study |
| pET28acsdA | pET28a plasmid for over-expression of N-terminal His tagged CSDA of *P. syringae*. | This study |
| pET28adbpa | pET28a plasmid for over-expression of N-terminal His tagged DBPA of *P. syringae*. | This study |
| pET28arhlB | pET28a plasmid for over-expression of N-terminal His tagged RHLB of *P. syringae*. | This study |

**Table S2.**
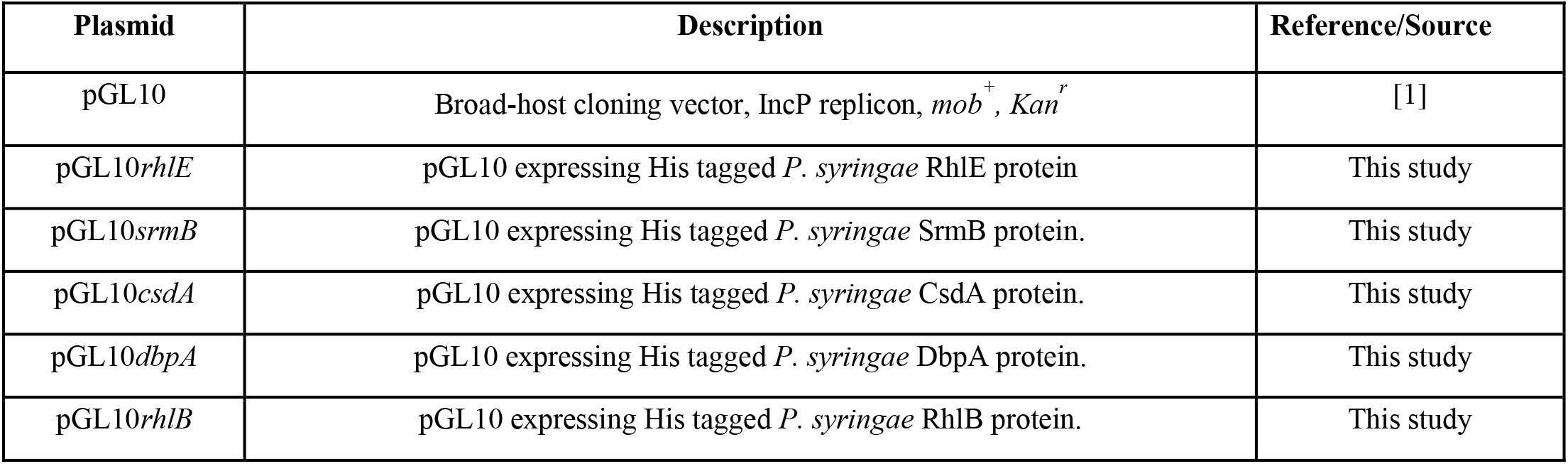
Plasmids used for functional complementation studies of RNA helicase mutants.

**Table S3.** Plasmids used as source of selective markers [antibiotic cassettes] and suicidal vector constructs for disruption of RNA helicase genes in *P. syringae*.

| Plasmid | Description | Reference/Source |
| --- | --- | --- |
| pTc28 | 2.4-kbp tet gene block from pOT182 cloned in pBluescript, Plasmid with Tetracycline resistance cassette | [2] |
| pKRP13 | pKRP13, Amp<sup>r</sup>, spc/str<sup>r</sup> Plasmid with spectinomycin resistance cassette. | [3] |
| pUC4K | Cloning vector, kan<sup>r</sup> Plasmid with Kanamycin resistance cassette | [4] |
| pJQ200SK | mob<sup>+</sup>, Gm<sup>r</sup>, Suicidal vector used for gene disruption | [5] |
| pJQrhIE-tet | 410 base pairs from 5' end-tet cassette-526 base pairs from 3' end of *rhIE* gene in pJQ200SK, Gm<sup>r</sup>, tet<sup>r</sup>; a suicidal construct for disruption of *rhIE* gene. | This study |
| pJQsrmB-kan | 422 base pairs from 5' end-kan cassette-519 base pairs from 3' end of *srmB* gene in pJQ200SK, Gm<sup>r</sup>, kan<sup>r</sup>, a suicidal construct for disruption of *srmB* gene. | This study |
| pJQcsdA-spec | 817 base pairs from 5' end-spec-cassette-857 base pairs from 3' end of *csdA* in pJQ200SK, Gm<sup>r</sup>, spec<sup>r</sup> a suicidal construct for disruption of *csdA* gene. | This study |
| pJQdbpA-spec | 710 base pairs from 5' end-spec cassette-676 base pairs from 3' end of *dbpA* gene in pJQ200SK, Gm<sup>r</sup>, spec<sup>r</sup>; a suicidal construct for disruption of *dbpA* gene. | This study |
| pJQrhIB-spec | 786 base pairs from 5' end-spec cassette-705 base pairs from 3' end of *rhIB* gene in pJQ200SK, Gm<sup>r</sup>, spec<sup>r</sup> a suicidal construct for disruption of *rhIB* gene. | This study |

**Table S4.**
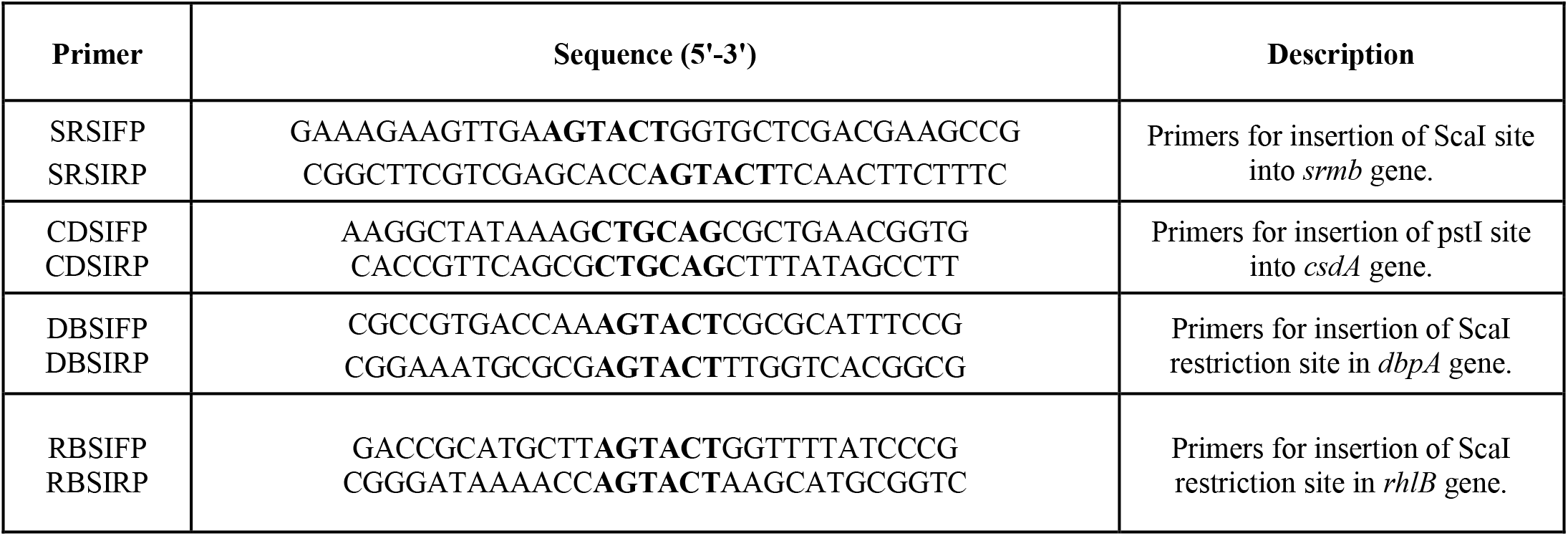
Primers used for insertion of restriction sites in RNA helicase genes to facilitate insertion of antibiotic resistance cassette.

**Table S5.**
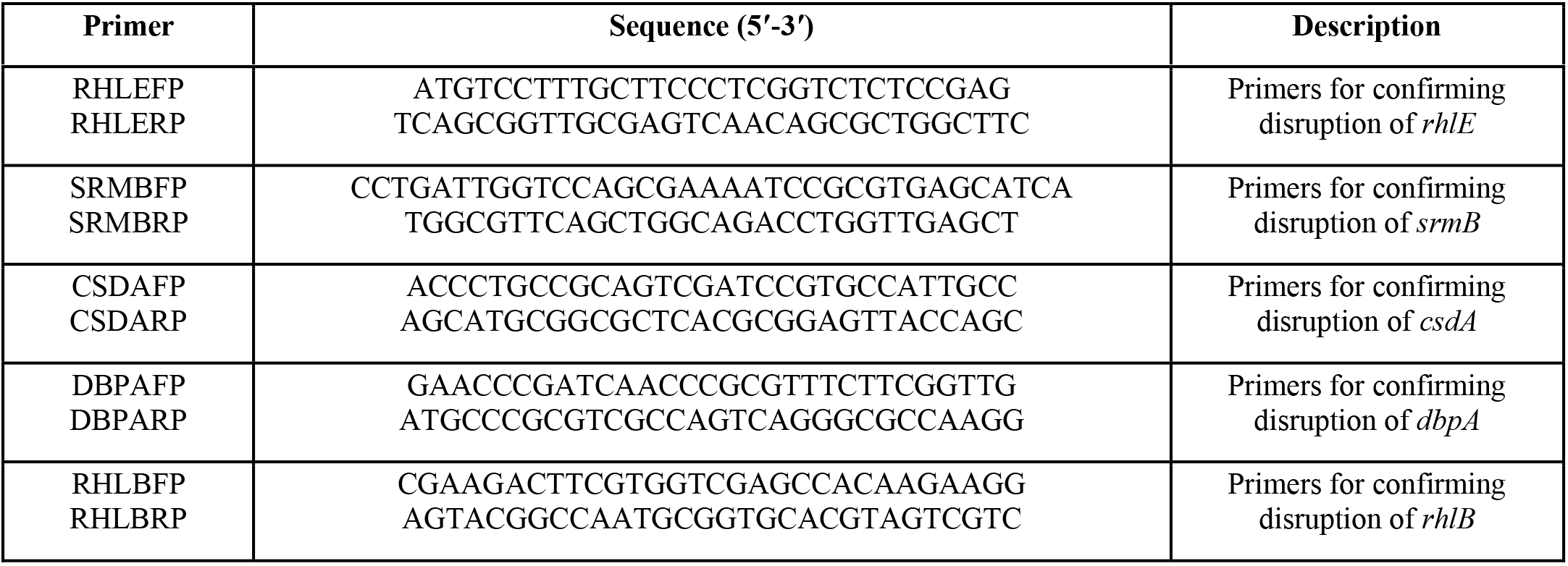
Primers used validation of RNA helicase gene disruption in *P. syringae*.

**Fig. S1.**
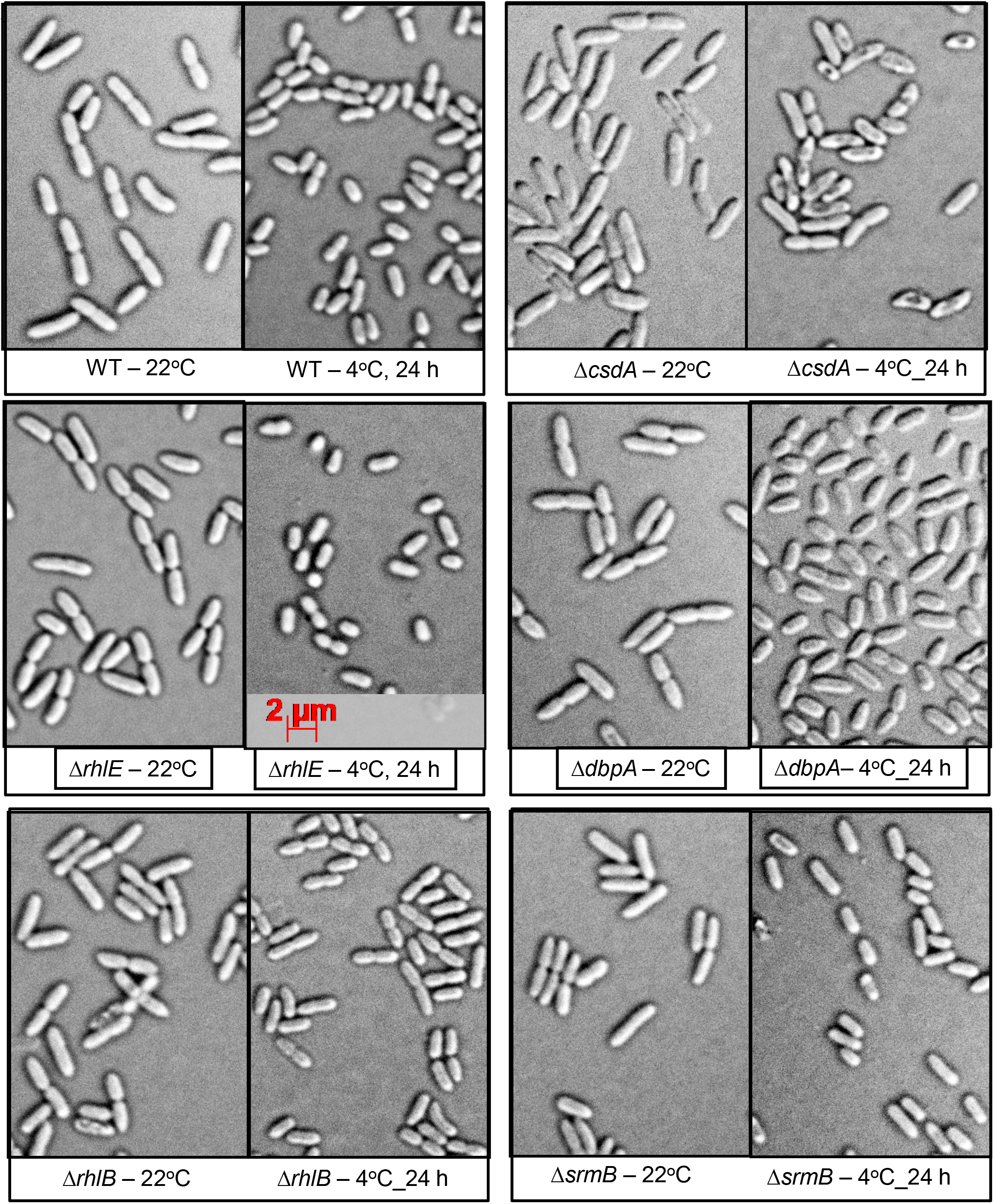
Morphological changes in RNA helicase mutants of *P. syringae* at 22 °C and 4 °C.

**Fig. S2.**
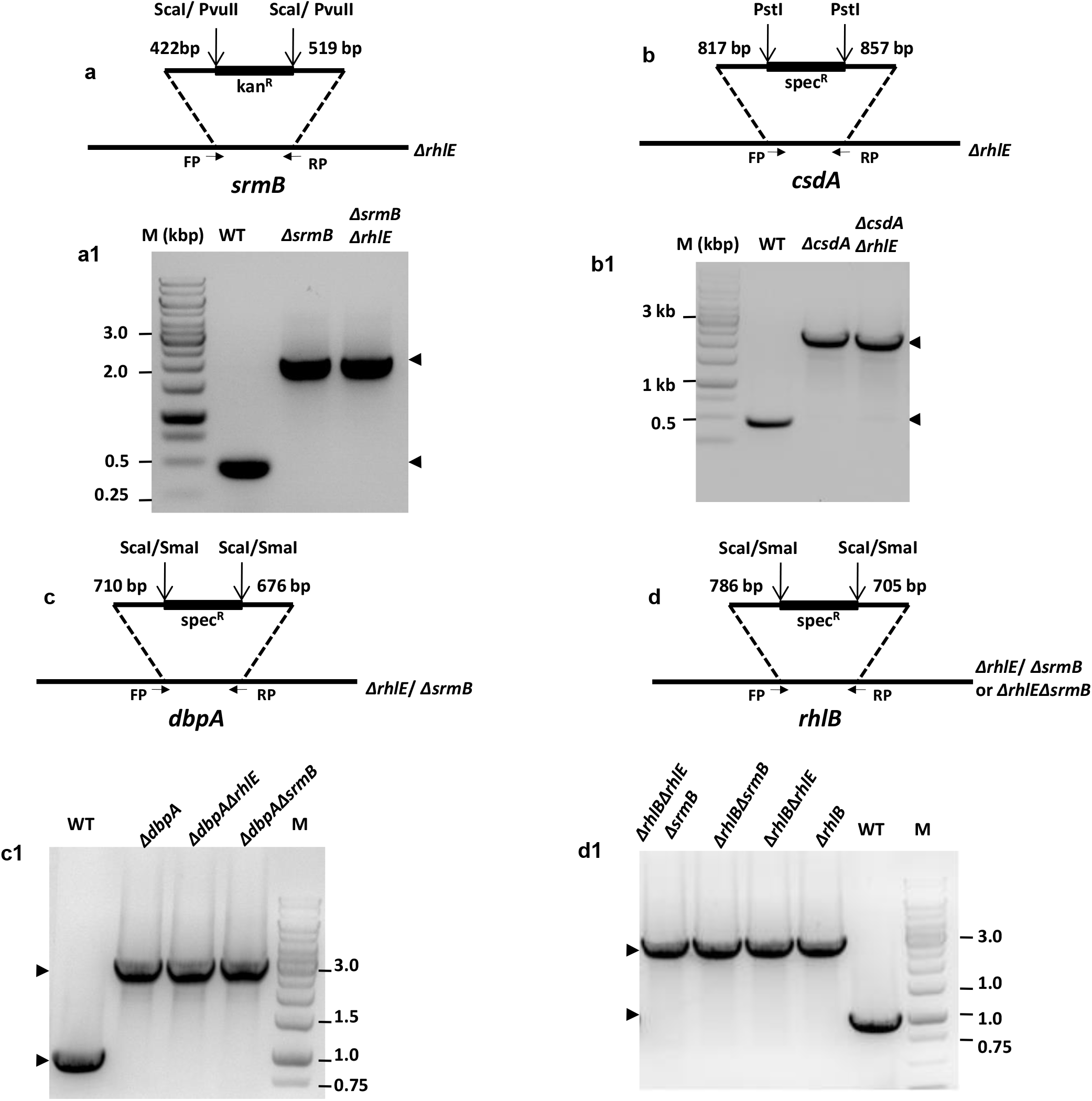
Recombinant vector construction and validation of mutants by PCR. (a) Schematic representation of suicidal construct pJQ*srmB-* kan used for disruption of *srmB* gene in wild type and Δ*rhlE* genetic backgrounds. (a1) Lanes marked as WT, Δ*srmB* and Δ*srmBΔrhlE* represent the PCR amplification results of strain specific gnomic DNA’s with *srmB* specific primers designed 225 base pairs from each end of inserted kan^r^ cassette. (b) Schematic representation of suicidal construct pJQ*csdA-* spec used for disruption of *csdA* gene in wild type and Δ*rhlE* genetic backgrounds. (b1) Lanes marked as WT, Δ*csdA* and Δ*csdAΔrhlE* represent the amplification results of strain specific gnomic DNA’s with *csdA* specific primers designed 250 base pairs from each end of spec^r^ cassette (c) Schematic representation of suicidal construct *pJQdbpA-spec* used for disruption of *dbpA* gene in wild type, Δ*rhlE* and Δ*srmB* genetic backgrounds. (c1) Lanes marked as WT, Δ*dbpA, ΔdbpAΔrhlE* and Δ*dbpAΔsrmB* represent the PCR amplification results of strain specific gnomic *DNA’s* with *dbpA* specific primers designed 500 base pairs from each end of inserted spec^r^ cassette. (d) Schematic representation of suicidal construct pJQ*rhlB-* spec used for deletion of *rhlB* gene in wild type, Δ*rhlE, ΔsrmB* and Δ*rhlEΔsrmB* genetic backgrounds. (d1) Lanes marked as Δ*rhlB, ΔrhlBΔrhlE, ΔrhlBΔsrmB* and Δ*rhlBΔrhlEΔsrmB* represent the PCR amplification results of Strain specific gnomic *DNA’s* with *rhlB* specific primers designed 500 base pairs from each end of inserted spec^r^ cassette.

## Construction of double and triple-deletion strains for helicases genes

The double and triple-deletion mutants were constructed by gene replacement of a second helicase gene in the first mutant background, using homologous recombination method [Fig S2]. However, we encountered limitation in the present study in the form of antibiotic resistance selection markers that were employed. We could use only three antibiotic resistance markers (*tet^R^, kan^R^* and *spec^R^*) which work well in *P. syringae* Lz4W, in terms of their expression and mutant selection on agar-plates, as shown for the single-deletion mutants, e.g., Δ*rhlE (rhlE::tet^R^), ΔrhlB (rhlB::spec^R^), ΔsrmB (srmB::kan^R^), ΔdbpA (dbpA::spec^R^*), and Δ*csdA (csdA::spec^R^*). As a result, for the combinatorial-deletion mutants we were successful in generating six double-deletion mutants (Δ*rhlEΔrhlB*), (Δ*rhlEΔsrmB*), (Δ*rhlEΔdbpA*), (Δ*rhlEΔcsdA*), (Δ*rhlBΔsrmB*), and (Δ*dbpA ΔsrmB*) out of ten possible double-deletion combinations (Δ*rhlEΔrhlB*, Δ*rhlEΔsrmB*, Δ*rhlEΔdbpA*, Δ*rhlEΔcsdA*, Δ*rhlBΔsrmB*, Δ*rhlBΔdbpA*, Δ*rhlBΔcsdA*, Δ*dbpAΔsrmB*, Δ*dbpAΔcsdA*, and Δ*csdAΔsrmB*). The three combinations (Δ*rhlBΔdbpA*), (Δ*rhlBΔcsdA*) and (Δ*dbpAΔcsdA*) were not obtained due to antibiotic selection difficulty, as the three genes (*rhlB, dbpA*, and *csdA*) had *spec^R^* disruption requiring common spectinomycin resistance selection for the two alleles. The other double-deletion mutant (Δ*csdAΔsrmB*)requiring *spec^R^* and *kan^R^* selection however repeatedly failed possibly due to lethality of this double mutant (see below). We also generated a triple-deletion mutant, namely (Δ*rhlEΔrhlBΔsrmB*), by successive selection against the three antibiotic resistance markers in the three genes (*rhlE::tet^R^), (rhlB::spec^R^*), and (*srmB::kan^R^*) respectively.

For the construction of (Δ*rhlEΔsrmB*), (Δ*rhlEΔcsdA*), (Δ*rhlE ΔdbpA*), and (Δ*rhlEΔrhlB*) doublemutants, the suicidal gene disruption plasmid vectors *pJQsrmB::kan^R^, pJQcsdA::spec^R^, pJQdbpA::spec^R^*, and pJQ *rhlB::spec^R^*, were mobilized individually from *E. coli* S17-1 strain into Δ*rhlE* mutant of *P. syringae* by biparental conjugation. The exconjugants were selected on ABM-agar plates containing ampicillin, tetracycline, and an additional appropriate antibiotic plus 5% sucrose. The additional antibiotics were kanamycin for the (Δ*rhlEΔsrmB*) selection, and spectinomycin for the other three, (Δ*rhlEΔrhlB), (ΔrhlEΔcsdA*) and (Δ*rhlEΔdbpA*) selections. Significantly, however, we failed to obtain the (Δ*csdAΔsrmB*) mutant requiring *spec^R^* and *kan^R^* selection repeatedly, in which the plasmid *pJQcsdA::spec^R^* was mobilized into Δ*srmB(kan^R^*) mutant background for the double deletion of *srmB* and *csdA* genes. We believe that the failure was not due to the synergistic adverse effects of ampicillin, tetracycline and kanamycin, simultaneously added to the selection medium, as we were able to select (Δ*rhlEΔsrmB*) mutant under the similar conditions. We surmise that the simultaneous removal of both SrmB and CsdA helicases, which are individually important during the low temperature (4°C) growth, is lethal to the cells even at the optimum temperature (22°C), as CsdA depleted cells (4°C lethal) grow poorly at 22°C.

The *dbpA* gene disruption in the double-mutant (Δ*rhlEΔdbpA*) was confirmed along with (Δ*dbpAΔsrmB*) mutant, both of which were created in parallel by mobilizing the *pJQdbpA::spec^R^* plasmid into Δ*rhlE* and Δ*srmB* mutant strains, respectively (Fig. 5.14). The (Δ*dbpAΔsrmB*) was selected by its resistance against the kanamycin and spectinomycin.

The double mutants (Δ*rhlEΔrhlB*) and (Δ*rhlBΔsrmB*) and the triple mutant (Δ*rhlBΔrhlE ΔsrmB*) were also constructed in parallel by mobilizing the pJQ*rhlB::spec^R^* plasmid into the parental mutant strains Δ*rhlE*, Δ*srmB*, and (Δ*rhlEΔsrmB*). The disruption of different RNA helicases in desired genetic backgrounds to generate double and triple helicase mutant strains was confirmed with PCR by using a set of gene specific primers. The mutants were also confirmed by western blot analysis [Data not shown].

## Notes

### Competing Interest Statement

The authors have declared no competing interest.

